# Stimulative piezoelectric nanofibrous scaffolds for enhanced small extracellular vesicle production in 3D cultures

**DOI:** 10.1101/2024.04.12.589114

**Authors:** James Johnston, Hyunsu Jeon, Yun Young Choi, Gaeun Kim, Tiger Shi, Courtney Khong, Hsueh-Chia Chang, Nosang Vincent Myung, Yichun Wang

## Abstract

Small extracellular vesicles (sEVs) have great promise as effective carriers for drug delivery. However, the challenges associated with the efficient production of sEVs hinder their clinical applications. Herein, we report a stimulative 3D culture platform for enhanced sEV production. The proposed platform consists of a piezoelectric nanofibrous scaffold (PES) coupled with acoustic stimulation to enhance sEV production of cells in a 3D biomimetic microenvironment. Combining cell stimulation with a 3D culture platform in this stimulative PES enables a 15.7-fold increase in the production rate per cell with minimal deviations in particle size and protein composition compared with standard 2D cultures. We find that the enhanced sEV production is attributable to the activation and upregulation of crucial sEV production steps through the synergistic effect of stimulation and the 3D microenvironment. Moreover, changes in cell morphology lead to cytoskeleton redistribution through cell–matrix interactions in the 3D cultures. This in turn facilitates intracellular EV trafficking, which impacts the production rate. Overall, our work provides a promising 3D cell culture platform based on piezoelectric biomaterials for enhanced sEV production. This platform is expected to accelerate the potential use of sEVs for drug delivery and broad biomedical applications.

## 1. Introduction

Extracellular vesicles (EVs) are nano/micro-sized lipid particles that are naturally secreted by most eukaryotic cells.^1^ Small extracellular vesicles (sEVs) with diameters of 50–200 nm play a critical role in facilitating intercellular communication,^2,3^ and have garnered significant interest as potential vehicles for drug delivery due to their numerous advantageous properties.^4–8^ Primarily, sEVs exhibit a robust capacity for the encapsulation and transference of bioactive molecules, including proteins, lipids, and nucleic acids. Furthermore, the inherently low immunogenic profile and high biocompatibility of sEVs mitigate the risk of eliciting adverse immunological responses, thereby enhancing patient safety upon administration. Moreover, sEVs possess the optimal dimensional and structural configuration required for carriers in drug delivery applications.^9^ Hence, there have been tremendous efforts to advance the development of methodologies for the scalable production, isolation, and functionalization of sEVs for clinical and therapeutic applications.

The traditional approach to producing EVs for biomedical applications involves the extraction of media from two-dimensional (2D) cell cultures.^10,11^ However, 2D cell cultures lack certain cell–cell and cell–matrix interactions, resulting in limited efficiency of sEV production.^12^ For instance, the standard production method through 2D cell culture systems produces 20–300 sEVs cell^-1^ hr^-1^ depending on the cell line, providing challenges for the efficient production of effective doses at 10^9^–10^11^ sEVs mL^-1^ on a daily time scale.^10^ To address this limitation, several strategies for stimulating the cells have been developed as a means of enhancing the production efficiency of 2D cultures. These strategies include chemical, pH, mechanical, electrical, electroporation, hypoxia, and gene expression strategies, as well as exposure to oxidative, thermal, or radiative stress.^13–24^ Reports suggest that mechanical and electrical stimuli can enhance EV production without affecting their size or cargo.^19,25^ These strategies use high magnitudes of electricity, or high frequency mechanical waves to induce EV production through manipulating the cell membrane structure, resulting in EV production enhancements of 1.7–2.1-fold hr^-1^.^25^ Despite the advances, these strategies can also lead to relatively low cell viability and the production of immunogenic EVs due to induced stress, posing challenges for their medical applications.^13,15^ Recently, three-dimensional (3D) cell cultures have been developed to improve EV production efficiency by ∼3-fold over that of the standard 2D Petri dish cultures.^26,27^ Biomaterial-based 3D culture platforms^26^ provide cells with a suitable 3D microenvironment through their biomimetic properties, such as porosity, pore size, and mechanical strength.^28–30^ Such a strategy can enhance EV production by up to three times and result in higher biomarker expressions on the produced EVs, indicating higher activity of EV biogenesis.^26^ Despite these advances in sEV production methods, sustainable production that meets the requirements of clinical applications remains challenging. Therefore, innovative biomanufacturing platforms for high-efficiency and high-quality sEV production have become a central focus in the field of biomedical science and engineering.

Nanofibrous scaffolds have been widely used for 3D cell cultures in tissue engineering due to their biomimetic properties, such as variable porosity, a high surface–volume ratio, and structural similarity to the extracellular matrix (ECM).^31–33^ One advantage is the integration of stimuli-responsive materials into nanofibrous structures. These materials can respond to various stimuli present in the tissue microenvironment, such as changes in temperature, pH, and mechanical forces, closely emulating the dynamic conditions of the ECM.^34–36^ By combining the effects of external cell stimuli with a 3D biomimetic microenvironment, these scaffolds enhance the cell– cell and cell–matrix interactions. Hence, stimulative nanofibrous scaffolds not only mimic the natural tissue settings more closely, but also have the potential to improve the efficiency and quality of sEV production.

In this study, we developed a tunable, stimulative 3D culture platform using piezoelectric nanofibrous scaffolds. This is the first 3D culture platform to achieve controlled piezoelectrical stimuli for enhanced sEV production (**Fig. 1A**).^37^ Piezoelectric polymers have recently been used to build nanofibrous scaffolds for energy storage, stimulatory cell cultures, and dynamic sensors.^38–40^ Specifically, piezoelectric nanofibers have been used in various biomedical applications such as tissue regeneration, where electrical stimulation causes cellular migration, and enhanced proliferation.^41,42^ In this regard, piezoelectric nanofibers provide controlled stimuli to cells by converting mechanical forces to electric potential through the direct piezoelectric effect.^41^ Piezoelectric nanofibers are easy to fabricate and can be finely tuned through the electrospinning process, offering precise control over their properties.^42^ Moreover, when used as tissue culture scaffolds, these piezoelectric nanofibers closely mimic certain bioelectrical properties by providing electrical stimuli through the cell–cell communication commonly found in natural cell microenvironments such as nervous tissue, liver tissue, and breast tissue.^43,44^ In this study, we fabricated the piezoelectric scaffolds (PESs) using polyacrylonitrile (PAN), and optimized the structural parameters for ideal 3D cell culture conditions by tuning the porosity, pore size, and thickness using gas foaming techniques.^45^ Our data demonstrates that the 3D cell culture in PES increases the production rate of sEVs per cell by a factor of 15.7 in HepG2 cells and by a factor of 6.7 in 3T3 cells, compared with 2D culture; importantly, the yielded sEVs are intact under the stimulated conditions. In addition, the PES platform demonstrates a method for safe, low voltage stimulation resulting in cell viability of over 90%. By investigating the underlying mechanism, we discovered that this significant enhancement in EV production is attributable to the activation and upregulation of crucial sEV production steps through the synergistic effect of stimulation and the 3D microenvironment. This is confirmed by a 1.5-fold rise in intracellular calcium ions and a 40% increase in metabolite concentration. Moreover, we find that the enhancement in sEV production is correlated with the cell morphology across different cell lines and culture conditions, potentially contributing to the cytoskeleton changes due to cell–matrix interactions in 3D cultures that facilitate intracellular EV trafficking. Overall, this study provides a promising platform for overcoming the limitations of EV production by improving the production rates and size distribution of EVs for drug delivery and broad biomedical applications.

**Fig. 1.**
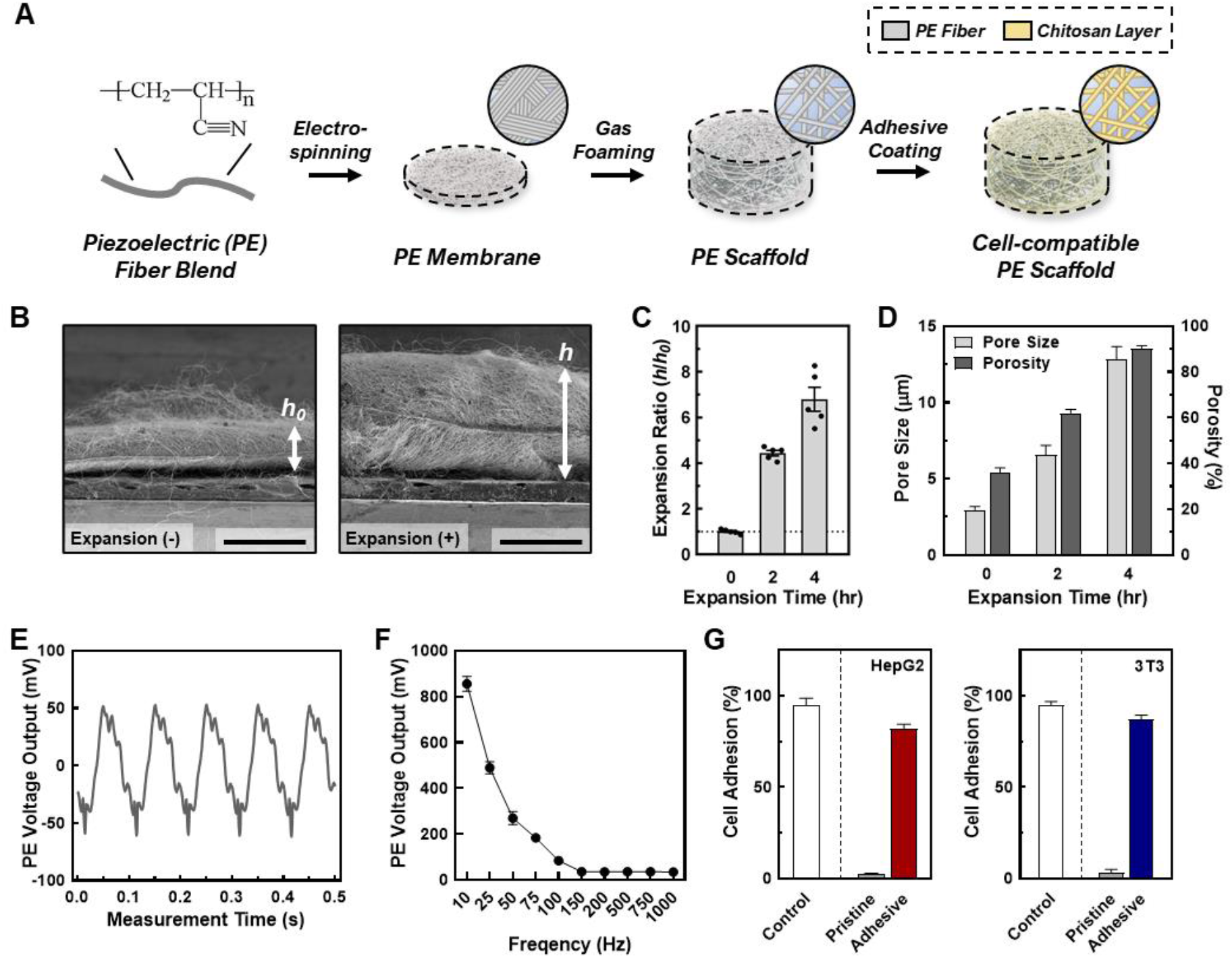
**(A)** A schematic for developing and processing the cell-compatible piezoelectric scaffold (PES). **(B)** Scanning electron microscopy (SEM) images of the piezoelectric scaffold cross section with and without gas foaming. **(C**–**D) (C)** The expansion ratio (*h/h*_*0*_) and **(D)** the pore size and porosity of PES with varying expansion times. Data shown as ± SD of 5 replicate samples. **(E**–**F)** the effect of the acoustic frequency on the piezoelectric properties of the scaffolds. Data shown as ± SD of 5 replicate samples. **(G)** The effect of chitosan coating of the scaffolds on the cells (Left: HepG2; Right: 3T3) seeding efficiency. Data shown as ± SD of 5 replicate samples.

## 2. Experimental

### 2.1 Materials and Reagents

Polyacrylonitrile (PAN; 181315), sodium borohydride (213462), and Pluronic f-127 powder (9003-11-6) were purchased from Sigma Aldrich (MA). N,N-dimethylformamide (DMF;D119-4) was purchased from Fischer Scientific (MA). Ethyl alcohol (3791-10), 99% acetic acid (BDH3092) and 0.22 μm vacuum filters (76010-388) were purchased from VWR (PA). Chitosan powder (c1569) was purchased from Spectrum Chemical (NJ).

Minimum Essential Medium (MEM; 10-010-CV) and Corning SpinX centrifuge filters (431491) were purchased from Corning (NY). Fetal bovine serum (FBS; 26140079), 100X antibiotic-antimycotic (15240062), 0.25% trypsin-EDTA (25200072), and Prestoblue (A13261) were purchased from ThermoFisher (MA). 4% paraformaldehyde in 0.1M phosphate buffer (15735) was purchased from Electron Microscopy Science (PA). Cell counting kit-8 (CCK-8;850-039-kl01) was purchased from Enzo Life Sciences (NY). 3D Celltiter-glo (G968A) was purchased from Promega (WI). Fura 2-AM (F1221) was purchased from Invitrogen Life Technologies (CA). HEPES buffered saline solution (C-40020) was purchased from PromoCell (Heidelberg, Germany). the LIVE/DEAD™ Cell Imaging Kit (488/570) was purchased from Thermofisher Scientific (MA).

### 2.2 Preparation of PAN solution and PAN nanofiber scaffolds

Solution property characteristics were performed similarly to prior work. ^42^ Solution viscosity was measured using a CPA-40 spindle connected to a Brookfield DV-I Prime viscometer (Brookfield, Toronto, Canada). The rotational speed of the spindle was ramped up from 0.5 rpm to whichever speed at which the torque reached closest to 100% (at least above 95%). After confirming that viscosity was independent of the shear rate, the viscosity value at maximum torque was recorded. Surface tension was measured using an automatic surface tensiometer (QBZY-1; Shanghai Fangrui Instrument, Shanghai, China), which had a platinum-coated plate connected to a hook. The force exerted on the hook as the plate came in contact with the solution was converted into surface tension values Arduino code (Atlas Scientific, NY) was used to take electrical conductivity measurement through a glass-body electrical conductivity probe (K = 0.1, Oakton) paired with an embedded conductivity circuit (EZO-EC; Atlas Scientific, NY) and an Arduino Uno Rev3 board. All solution property measurements were taken at room temperature immediately before or after electrospinning to correlate them most closely with the resulting nanofiber properties.

Nanofibers with a diameter of 760 nm were produced through an electrospinning process. A solution of 10 wt% PAN was prepared in DMF. Electrospinning was carried out under specific conditions, namely an electrospinning distance of 10 cm, an applied voltage of 13 kV, and a solution feed rate of 1 mL hr^-1^. This process was conducted in a controlled environment of 23°C and 40% relative humidity. The resulting nanofibers were collected on a rotating collector drum covered with aluminum foil, operating at 400 rpm. The electrospinning duration was optimized to achieve nanofibers with the desired thickness of approximately 100 μm.

### 2.3 Gas foaming expansion of PAN scaffolds

Gas foaming expansion of the PAN scaffolds was performed following a previous study.^45^ A 2 cm × 2 cm × 100 μm scaffold was submerged in a freshly prepared 1M sodium borohydride solution for varying lengths of time (0, 2, and 4 hr) at 22°C. The expanded nanofiber scaffolds were gently transferred into a separate beaker and washed three times with DDI water before being freeze-dried for 24 hr. The sample thickness was measured using a ruler and the morphology was documented via a digital camera.

### 2.4 Piezoelectric property measurement

Nanofiber scaffolds were prepared in a cantilever setup, similar to prior work.^42^ This setup allows for the controlled application of strain to the samples while simultaneously measuring their electrical output. The PAN nanofiber scaffolds were cut into strips of size 4 × 1.2 cm, and brass slabs of size 7.2 × 1.6 × 0.01 cm^3^, electrically isolated with polyimide tape, were employed as electrodes to measure the voltage. One brass slab was in direct contact with the nanofiber sample, secured with double-sided copper tape, while the other slab remained unexposed. Two 24-gauge wires were soldered to these electrodes, sealed with polyimide tape, and connected to a breadboard with inputs to a PicoScope 2204A (Pico Technology Ltd., Cambridgeshire, UK) for voltage measurement. To induce controlled strain, a 2.3 g proof mass was placed on the cantilever’s end, driven by a custom-made oscillatory system, with the cantilever holder clamped atop a subwoofer diaphragm. The strain was calculated using the formula:

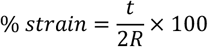

with *t* representing the cantilever thickness and *R* being the radius of curvature, as determined through a surface-mounted camera. A sinusoidal sine wave with a controlled amplitude and a 10 Hz frequency was applied to the speaker system, and the voltage output was measured.

### 2.5 Scanning electron microscopy images of scaffolds

Scanning electron microscopy (SEM; Thermo Prima environmental-SEM; ThermoFisher, MA) was used to image samples with and without cells. Cell-free samples were fixed on a metallic stud (75210; Electron Microscopy Science, PA) with double-sided conductive tape and sputter-coated with gold before imaging under 10 kV. The sample thickness, average fiber diameter, and pore size were measured based on the SEM images using the ImageJ software. For the SEM imaging of cells seeded on the nanofiber scaffold, the cells were fixed with 2% paraformaldehyde for 1 hr. Samples were then washed three times with PBS and dehydrated sequentially in 50% ethanol (2 × 10 min), 70% ethanol (2 × 10 min), 80% ethanol (2 × 10 min), 95% ethanol (2 × 10 min), and 100% ethanol (3 × 10 min). Samples were then dried at room temperature and fixed on a metallic stud with double-sided conductive tape before being imaged. SEM images of cells were processed and measured for roundness, axial ratio, and cell area using the default measurement programming in ImageJ, as described in previous work.^46^ The following equations were used in measuring the cell morphology:

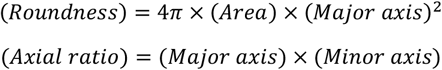

These measurements were used to quantitatively compare the cell morphology between 2D culture, PES OFF, and PES ON samples.

### 2.6 Porosity measurements of PAN scaffolds

The porosity of the nanofiber scaffolds was calculated according to the liquid displacement of each sample. The mass of each sample was measured before and after being submerged in water. The porosity was calculated using the following equation:

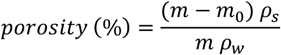

where *m*_0_ and *m* are the masses before and after being submerged in water, respectively, while *ρ*_*w*_ and *ρ*_*s*_ are the densities of water and the PAN bulk material, respectively.

### 2.7 2D cell culture and media collection

Human hepatocellular carcinoma cell line (HepG2) and mouse embryonic fibroblast cell line (3T3) were cultured separately on six well plates at a density of 3 × 10^5^ cells/well in MEM containing 10% FBS and 1% 100 X Antibiotic-Antimycotic. The medium was exchanged every 48 hr. Once the culture reached 70% confluency, the serum-containing medium was replaced with serum-free medium and incubated for 24 hr before being collected for sEV isolation. Cells were then harvested after media collection with Trypsin-EDTA for 5 min followed by spinning down the cells at 1000 rpm for 5 min. The cells were resuspended in fresh MEM and diluted appropriately to be counted using a hemocytometer.

### 2.8 3D cell culture on PAN scaffolds and media collection

The scaffolds were cut into 4 cm × 2 cm strips and prepared for seeding, which included washing, coating with chitosan, and sterilization. Each sample was rinsed in distilled water for 30 min and transferred to a solution of 1 mg mL^-1^ chitosan in 0.1 M acetic acid for 30 min. The coated nanofiber scaffolds were then rinsed with fresh distilled water for 30 min and air dried. The processed samples were placed in a 60 mL petri dish and UV-sterilized before seeding. HepG2 and 3T3 cells were passaged and seeded separately at 3 × 10^5^ cell scaffold^-1^. The seeded scaffolds were then incubated for 30 min at 37°C before the addition of serum-containing MEM and incubation at 37°C and 5% CO_2_. After 48 hr, the conditioned medium was replaced with serum-free medium and incubated for 24 h before being collected for sEV isolation. Cells were then harvested after media collection with Trypsin-EDTA for 5 min followed by spinning down the cells at 1000 rpm for 5 min. The cells were resuspended in fresh MEM and diluted appropriately to be counted using a hemocytometer. For the Live/Dead assay, the LIVE/DEAD™ Cell Imaging Kit was used following the manufacturer’s instructions. For the staining, each scaffold whether embedding cells were exposed to 1 mL of final staining solution which was a 1:2 mixture of staining agent from the product and fresh culture media.

### 2.9 Acoustic stimulation and media collection

3D culture samples containing serum-free medium were stimulated using sinusoidal acoustic waves (3-Ω subwoofer; PS-EW1-2; Samsung Electronics, Suwon-si, Republic of Korea) in a sound-controlled box. Samples were stimulated for 15 min at an amplitude of 85 dB and frequency of 100 Hz. The samples were incubated at 37°C and 5% CO_2_ for 24 hr before being collected for sEV isolation. The cells were then detached and counted, similar to 2D and 3D cultured samples.

### 2.10 Cell seeding efficiency measurements for nanofiber scaffolds

HepG2 and 3T3 cells were seeded separately on 2 mm × 2 mm scaffolds (*N=*3) with and without a chitosan coating at a density of 5 x 10^5^ cells scaffold^-1^ and incubated in serum-containing MEM at 37°C and 5% CO_2_ for 24 hr. The cell seeding efficiency was measured in two different tests. One test calculated the number of cells in the scaffold, and the other test calculated the number of cells out of the scaffold:

1. The cultured medium was aspirated, and the scaffolds were washed with PBS thrice before the addition of 10% CCK-8 reagent in cell culture media and incubation for 4 hr at 37°C and 5% CO_2_. For Pesto-blue assay, standard protocol from vendor was applied. The cell count was then measured through absorbance at 460 nm wavelength, and the seeding efficiency was calculated using the following equation:

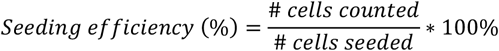 The seeding efficiency was calculated for scaffolds with and without chitosan coating, and for monolayer cultures.
2. The cell medium during each step of EV production was collected and measured for cell count using the above CCK-8 method.

For the confocal imaging, the samples were kept in 2 mL PBS before imaging with A1R-MP Laser Scanning Confocal Microscopy (CLSM; Nikon, Tokyo, Japan).

### 2.11 Cell metabolism assay

CellTiter-Glo^®^ and CCK-8 assays were used to measure the cell metabolism of 2D and 3D cultured samples. Both assays were performed using the manufacturer’s protocol. **CellTiter-Glo**^**®**^: HepG2 and 3T3 were seeded on 2D and 3D culture samples in 96 well plates at different cell densities (1 × 10^4^, 2 × 10^4^, 4 × 10^4^, 8 × 10^4^, and 16 × 10^4^ cells sample^-1^) and incubated with 50 μL of serum-containing MEM at 37°C and 5% CO_2_ for 24 h. The medium was then removed, and the cells were washed twice with PBS before new medium was added. The samples were then equilibrated to room temperature for 30 min followed by the addition of 50 μL CellTiter-Glo^®^ 3D reagent to each sample. The samples were mixed vigorously for 5 min to induce cell lysis and allowed to incubate at room temperature for 25 min to stabilize the luminescent signal. The ATP levels were measured through luminescence measurements. **CCK-8:** The seeding, incubation, and medium change was the same as for the CellTiter-Glo^®^ assay. The CCK-8 working reagent was prepared by diluting 10 μL of CCK-8 stock reagent in 190 μL MEM. Then, 200 μL of the working reagent was added to each well and incubated at 37°C and 5% CO_2_ for 4 h. The cell metabolism was then measured through absorbance at 460 nm wavelength.

### 2.12 sEV isolation and concentration measurement

sEVs were isolated from the media via size-based separation. The isolated media first went through a 0.22 μm filter to capture larger vesicles and cell debris. The flowthrough solution was then added to a 100 kDa centrifuge filter and centrifuged at 200 g for 4 × 30 min time periods; the mixture was washed with PBS between each centrifuge session. The sEV solution was then concentrated down to 1 mL for sEV characterization. The sEV concentration and size distribution were measured through nanoparticle tracking analysis (NTA; Nanosight NS300; Malvern, Worcestershire, UK). Samples were diluted appropriately to maintain accurate particle counts. For each sample, five 60-sec videos were acquired at a camera level of 8 and detection threshold of 2. The laser chamber was cleaned with milliQ water between each sample reading to ensure no sample contamination occurred. The videos were analyzed using the NTA3.0 software to obtain the particle concentration, along with the mean and mode particle sizes of each sample.

### 2.13 Western blot

sEVs were lysed with 1 X RIPA buffer (9806; Cell Signaling Technology, USA), and the total protein concentration was quantified using Pierce BCA Protein Assay Kits (23225; Thermo Fisher, USA). The protein amount in lysed sEVs was estimated based on a calibration curve plotted by albumin (BSA) standards. Next, 12 μg of proteins from sEV lysates were denatured and loaded on sodium dodecyl-sulfate polyacrylamide gel electrophoresis (SDS-PAGE). The separated proteins were subsequently transferred onto a nitrocellulose membrane (1662807; Bio-Rad, USA) and blotted overnight with primary antibodies purchased from Santa Cruz Biotechnology, USA: anti-CD9 antibody (C-4), anti-CD63 antibody (MX-49.129.5), anti-CD81 antibody (B-11), anti-CD9 antibody, anti-HSP70 antibody (W27), anti-HSP90 antibody (F-8), and anti-beta-actin antibody (sc-47778). The secondary antibodies (anti-mouse HRP-linked antibody, 7076; Cell Signaling Technology, USA) were then treated for blotting and the HRP on the immunoblots was detected by Clarity Max Western Enhanced Chemiluminescence (ECL) Substrate (1705060; Bio-Rad, USA) using a ChemiDoc XRS+ System (Bio-Rad, USA). The relative expression levels of the detected proteins were quantified using the ImageJ software.

### 2.14 sEV production rate measurement using ExoELISA

An enzyme-linked immunosorbent assay (ELISA) of sEV solutions was performed using ExoELISA (System Biosciences, CA) following the manufacturer’s protocol. The sEV samples were prepared by adding 60 μL of sample and 60 μL coating buffer into triplicate wells. Standards were prepared using the manufacturer’s protocol. The samples were then incubated at 37°C for 1 hr, followed by washing with 1 X wash buffer three times for 5 min. The samples were then incubated with CD63 primary antibody at 37°C for 1 h followed by washing with 1 X wash buffer three times for 5 min. The samples were then incubated with CD63 secondary antibody at 37°C for 1 hr and washed with 1 X buffer three times for 5 min. Finally, the TMB ELISA substrate was added to the sample and incubated at room temperature for 15 min while shaking. After shaking, a stop buffer was added, and the absorbance was measured at 450 nm using a microplate reader (30190087; TECAN, Mannedorf, Switzerland).

### 2.15 Transmission electron microscopy of sEVs

The sEV solutions were negatively stained and imaged through transmission electron microscopy (TEM) using the Talios F200i (S)TEM (ThermoFisher, MA) at an 80 kV accelerating voltage. The TEM samples were prepared by adding 5 μL of 1 × 10^8^ particles mL^-1^ sEV solution to an ultrathin carbon film copper grid and incubating at room temperature for 2 min. The solution was then aspirated using filter paper and washed with 5 μL filtered distilled water for 10 sec. After aspirating the distilled water, 5 μL of Uranyless negative staining solution (22409; Electron Microscopy Science, PA) was added to the sample grid (CF200-CU-25; Electron Microscopy Science, PA) and incubated for 1 min. The Uranyless solution was aspirated, and the grid was left to dry before imaging.

### 2.16 Intracellular Ca^2+^ measurements

The intracellular Ca^2+^ concentration was measured using Fura 2-am following a similar protocol as in previous studies.^19,25^ Cells were seeded at a density of 3 × 10^5^ cells well^-1^ in a 6-well plate and incubated overnight at 37°C and 5% CO_2_. After incubation, they were treated with 10 μM Fura-2 AM in HEPES-buffered saline solution and pluronic f-127 for 1 hr in a humidified incubator. The cells were then washed to remove the extracellular dye and replenished with MEM. The appropriate samples were then exposed to acoustic stimulation and the changes in the fluorescence intensity were measured with a spectrophotometric plate reader (TECAN, Mannedorf, Switzerland).

### 2.17 Statistical analysis

The data presented in this paper are expressed as the mean +/-the standard error of replicate measurements and analyzed using one way ANOVA test, where applicable.

### 2.18 Cell-Free DNA (cfDNA) Isolation

cfDNA was extracted and isolated from concentrated sEVs from Experimental section 2.12 using Plasma/Serum Cell-Free Circulating DNA Purification Kit – Mini (55100; Norgen Biotek, Canada) per manufacturer instructions. The volume of concentrated sEVs used extraction were 500, 435, 145, 500, 280, and 85 μL for HepG2 2D-culture, HepG2 3D-culture without stimulation, HepG2 3D-culture with stimulation, 3T3 2D-culture, 3T3 3D-culture without stimulation, and 3T3 3D-culture with stimulation, respectively. A blank control sample was extracted in parallel using 500 μL of DNA Dilution Buffer (4405587C; Thermo Fisher, USA). All samples were diluted to 500 μL prior to extraction. All samples were each eluted into 30 μL of purified cfDNA.

### 2.19 TP53 Nested PCR and NRAS PCR of cfDNA

A wild-type sequence and associated primer sets within TP53 were previously reported.^47^ The TP53 nested PCR was performed using the outer and inner primer sets listed in **Table S1**. A wild-type sequence within NRAS was identified using NCBI Sequence Viewer for Homo sapiens chromosome 1, GRCh38.p14 Primary Assembly.^48^ Primer sequences were designed using Primer-BLAST service.^49^ PCR assays were designed for 20 μL reactions containing 10 μL SsoAdvanced™ Universal SYBR® Green Supermix (1725271; Bio-Rad, USA) and 200-nM of the appropriate forward and reverse primers (Integrated DNA Technologies, USA). 2 μL of the isolated cfDNA for each sample and the blank control were used as the template for the TP53 Outer and NRAS PCR reactions. 2 μL of the amplified PCR products of the TP53 Outer reactions were used as templates for the TP53 Inner PCR reactions. Cycling conditions for the TP53 Outer reactions were: 10 min at 95°C followed by 40-cycles of [30 sec at 95°C, 30 sec at 53°C, 1 min at 60°C] and ending with 2 min at 60°C. Cycling conditions for the TP53 Inner reactions were: 10 min at 95°C followed by 40 cycles of [30 sec at 95°C, 30 sec at 52°C, 1 min at 60°C] and ending with 2 min at 60°C. Cycling conditions for the NRAS reactions were: 2 min at 95°C followed by 40 cycles of [30 sec at 95°C, 20 sec at 50°C, 40 sec at 60°C] and ending with 2 min at 60°C. Amplified products were stored in 4°C until examination via gel electrophoresis.

### 2.20 cfDNA Gel Electrophoresis

Amplified PCR products were examined in a 1.5% Agarose-1 gel formulated using 1 X TAE Buffer (J63931.K2; Thermo Fisher, USA). Sample mixtures of 6 μL containing 1 μL of PCR product or GeneRuler 100 bp DNA Ladder (SM0243; Thermo Fisher, USA), 1 μL of DNA Gel Loading Dye (R0611; Thermo Fisher, USA), and 4 μL of DNA Dilution Buffer (4405587C; Thermo Fisher, USA) were loaded into each lane. Electrophoresis was run in 1 X TAE Buffer at 80 V for 70 min on a PowerPac™ Basic Power Supply (1645050; Thermo Fisher, USA). Upon completion, gels were removed from the electrophoresis unit and incubated away from light in 5 μL Thiazole Green, 10,000X (40086; Biotium, USA) diluted in 50 mL 1 X TAE Buffer for 30 min. Gels were examined using a blue-light transilluminator. Images were taken with a smartphone camera and processed using ImageJ.

## 3. Results and Discussion

### 3.1 Scaffold fabrication and processing for 3D cell cultures

For piezoelectric scaffold, PAN was selected due to its electrospinnable feature enabling the nanofibrous matrix, hydrophilic nature allowing 3D cell culture, and piezoelectric characteristics enabling the acoustic stimulation to cells. The PESs were first fabricated through electrospinning using 10 wt% PAN in DMF (*e*.*g*., Solution characteristic summarized in **Table S2**). Following successful fabrication, we tuned the scaffold parameters, including the pore size, porosity, and thickness, to ensure cell penetration and ample void space for cell growth.^50^ This was done by employing a gas foaming technique developed in our previous work.^45^ We examined the expansion effect of the gas foaming technique on the scaffold through SEM imaging (**Fig. 1B–1C** and **Fig S1**). During the gas forming processes, there was loosen porous structure observed (**Fig. S1A**) while no significant change in the average fiber diameter (pristine: 442 ± 39 nm, 2 hr: 459 ± 63 nm, 4 hr: 441 ± 83 nm) (**Fig. S1B**). The expansion ratio (*i*.*e*., *h*/*h*_*0*_ where *h*: expanded height of scaffold, *h*_*0*_: original height of scaffold) increased by a factor of 4.4 ± 0.3 after 2 hr and a factor of 6.8 ± 0.6 after 4 hr, similar to the results of previous studies (**Fig. 1D**).^45,51^ To ensure the scaffolds had the optimal structural parameters for cell seeding and cell growth, we measured the pore size and porosity of as-synthesized (*i*.*e*., 0 hr-expansion sample) and gas-foamed scaffolds. The gas foaming process resulted in scaffolds with significantly larger pore sizes (pristine: 3.0 ± 0.3 μm, 2 hr: 5.3 ± 0.8 μm, 4 hr: 12.6 ± 2.3 μm). Pore sizes above 10 μm facilitate enhanced cell penetration through the scaffold, ensuring an even cell distribution.^50^ To validate the porosity of the scaffolds, we conducted a water-displacement test on both pristine and processed fibrous scaffolds. The gas-foamed fibrous scaffolds exhibited ∼2.5-fold higher porosity in comparison with the pristine ones (pristine: 37.0 ± 6.9%, 2 hr: 62.3 ± 5.6%, 4 hr: 91.3 ± 3.7%), signifying greater void space for cell growth within the scaffolds. In addition, there was no observed change in the average fiber diameter after gas foaming (pristine: 442 ± 39 nm, 2 hr: 459 ± 63 nm, 4 hr: 441 ± 83 nm) (See **Fig. S1B**). Therefore, we used the PESs processed by gas foaming for 4 h for the subsequent work to assess sEV production in the 3D stimulative culture platform.

To verify whether the piezoelectric properties of the processed scaffolds were sufficient for inducing cell stimulation, we conducted a cantilever test to measure the mechanical strain of the fibers in conjunction with the electrical output (**Fig. 1E**). We performed the cantilever tests after the post-processing phase of scaffold fabrication and in a wet phase to mimic the scaffold structure during cell culture. In this test, we observed a peak-to-peak voltage output of 110 ± 15 mV, which is within the desired voltage output for facilitating safe cell stimulation.^52,53^ The effect of the acoustic frequency on the piezoelectric output (**Fig. 1F** and **Fig. S2**) was also investigated to optimize the conditions for cell stimulation. We found an inverse relationship between the acoustic frequency and piezoelectric output, with an active frequency range of 10–100 Hz for PESs. This was supported by our finding that PESs activated by acoustic frequencies above 100 Hz did not exhibit significantly different voltages or percentage strains. Additionally, acoustic frequency had no significant effect on the pore size of the scaffold, demonstrating the physical stability of our system for cell stimulation in 3D cell culture (**Fig. S3**). Based on these findings, we chose to use an acoustic frequency of 100 Hz to provide a sufficient voltage output in the PESs, from a strain of 0.01% produced by an acoustic amplitude of 85 dB.

To ensure that the cells adhere to the scaffolds, we coated the scaffolds with chitosan, a well-known bioactive polymer used in promoting cell adhesion, cell proliferation, and antibacterial properties.^54–57^ The chitosan coating alleviated the piezoelectric output from the scaffold without completely insulating its piezoelectric properties (**Fig. S4**). We tested the cell seeding efficiency of coated scaffolds, compared with unfunctionalized scaffolds (**Fig. 1G**; A photo of scaffolds before cell seeding is represented in **Fig. S5**). We chose two cell lines, a human hepatocellular carcinoma line (HepG2) and a mouse derived fibroblast cell (3T3), as proof of concept for the following cell related experiments. We found that the seeding efficiency for HepG2 on the chitosan-functionalized scaffolds (> 95%) was 25-fold higher than that of unfunctionalized scaffolds (3.2 ± 1.0%). This is due to the electrostatic interactions between the chitosan and the cell surface.^58^ Similar to the HepG2, we found a 26-fold increase in the seeding efficiency of 3T3 cells on the chitosan-functionalized scaffolds (functionalized: 87.4 ± 2.0%, unfunctionalized: 3.3 ± 1.7%), confirming sufficient cell adhesion to the functionalized scaffolds. Cell adhesion was also tested indirectly by tracking the number of cells dislodged from the scaffold. In doing so, we observed 88.3 ± 1.6% of cells adhered to the scaffold, supporting the enhanced cell adhesion capability of the chitosan coated PESs (**Table S3**). The cell adhesion to the scaffold was then imaged using confocal microscopy, in which cells were observed to be properly adhered (**Fig. S6**). As shown in the confocal microscopic imaging and its analysis, live and dead cells for both HepG2 and 3T3 cases (*e*.*g*., Green channel: Live cells; Red channel: Dead cells) were stably adhered to chitosan-coated scaffolds (**Fig. S6A**) throughout 3D spaces (**Fig. S6B**), demonstrating successful cell seeding on the scaffold and inside the scaffold. Besides, the chitosan coated scaffold showed great biocompatibility for both cell lines (*e*.*g*., > 80% for both 3T3 and HepG2; **Fig. S6C**). These results indicate that cells adhere to the PES with good biocompatibility, aligning with our previous tests on cell viability and adhesion efficiency. To confirm growth of cells on the scaffolds, we tested the proliferation of each cell line on the scaffolds over 13 days (**Fig. S7A**). We observed a 6.7 (± 0.6)-fold expansion in HepG2 cells and 7.7 (± 0.9)-fold expansion in 3T3 cells over 13 days. Therefore, we concluded that the cells can effectively reproduce on the scaffolds. We also observed a plateau in the cell count starting after 9 days of cell culture, indicating that the stationary phase of cell growth on the scaffold starts at 9 days. To ensure that activating the PES does not affect the cultured cells, we tested the cell viability at acoustic amplitudes ranging from 50– 110 dB with a frequency of 100 Hz and found that the cell viability remained above 93% under all conditions (**Fig. S7B**–**C**). Overall, the PESs with functionalization and activation exhibited high biocompatibility of both cell lines.

### 3.2 Synergistic effect of 3D culture and cell stimulation in PES on EV production

Upon successful cell seeding and culturing in the optimized PAN PESs, we tested the synergistic effect of the 3D culture and cell stimulation on sEV production (**Fig. 2A**). To confirm the quality of the sEVs, we analyzed their size, integrity, and morphology using NTA and TEM. We analyzed the size distributions of sEVs from both cell lines based on the NTA results (**Fig. 2B–C**; Raw particle count including fresh media shown in **Fig. S8**). According to NTA, cells cultured in all groups, including 2D culture (control), 3D culture in PES with (PES ON) and without stimulation (PES OFF), generated particles within the size range of sEVs,^1^ and the mean particle sizes among all groups were not significantly different (p > 0.05). The sEVs produced from HepG2 cells in the control group exhibited a mean particle size of 152 ± 38 nm, and average sizes of 136 ± 36 nm and 132 ± 40 nm in the PES OFF and PES ON groups, respectively (**Fig. S9**). The 3T3 cells exhibited particles with similar sizes among all groups (control: 141 ± 40 nm, PES OFF: 131± 32 nm, PES ON: 136 ± 37 nm). TEM images of the sEVs confirmed their stability, showing that sEVs in all groups were intact, round, and oval-like (**Fig. S10A**), with a narrow size distribution (**Fig. S10B**) without any significant distribution differences from NTA analysis.

**Fig. 2.**
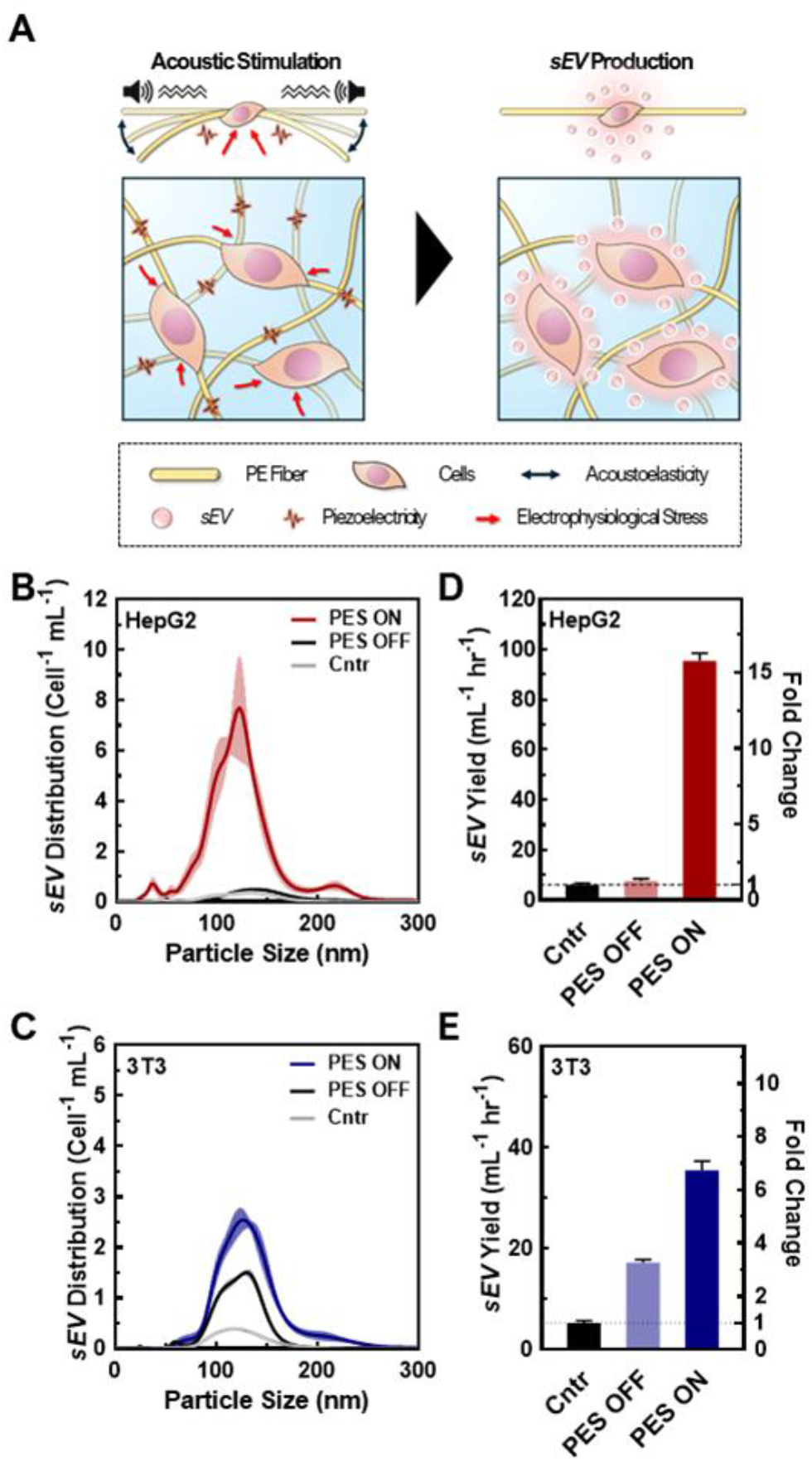
**(A)** Schematic of the piezoelectric stimulation that enhances sEV production. **(B**–**C)** Size distribution of sEVs derived from **(B)** HepG2 and **(C)** 3T3 cells using NTA. Data shown as ± SD of 5 replicate samples. **(D**–**E)** Measured EV production rate per cell from **(B)** HepG2 and **(C)** 3T3 using ExoELISA. Data shown as ± SD of 5 replicate samples.

To evaluate the production efficiency of sEVs from the PES ON group compared with the control and PES OFF groups, we assessed the sEV production with and without acoustic stimulation by measuring the sEV yield and the production rate per cell using the CD63 ELISA kit, ExoELISA (**Figs. 2D–E**). We found that HepG2 cells in the PES ON group produced sEVs with a 15.4-fold increase in yield and a 15.7-fold increase in production rate compared with the control (2D culture). We further tested the effect of cell stimulation on non-cancer cells using 3T3 cells, a fibroblast cell line, and found a similar trend. The 3T3 cells stimulated in the PES produced the largest yield and at the highest rate among all conditions. Conversely, the stimulation was not as effective on the EV production rate in the 3T3 cells (6.7-fold increase) as in the HepG2 cells (15.7-fold increase). These results agree with analysis from NTA measurements (**Fig. S11**). Additionally, the sEV distribution curves from control group with acoustic stimulation (*i*.*e*., Cntr ON) confirmed the increase in sEV yield was not induced by the acoustic stimulation only (**Fig. S12**).

To further verify the presence of sEVs produced from the PES platform, a western blot was conducted for sEV biomarkers (CD63, CD9, and CD81) (**Fig. 3A** and **Fig. S13**–**14**). The western blot results of PES-derived sEVs clearly displayed distinct bands for all sEV biomarkers, confirming the presence of sEVs from both HepG2 and 3T3 cell lines. To note, sample concentrations have been normalized by protein content and thus the level of sEV markers, CD9, CD63, and CD81 in WB are not informative regarding sEV quantity. We further compared the stress levels of sEVs across all groups (2D, PES OFF, and PES ON) based on the expression of heat shock proteins (HSPs) (HSP70: **Fig. 3B C**; HSP90: **Fig. 3D–E**). The relative expression levels were normalized against β-actin, which was used as the loading control in this western blot analysis. While there was no significant difference in the expression level of HSP70 among all groups within both cell lines, distinctions in HSP90 levels were observed. The PES OFF group exhibited the highest level of HSP90 expression, while the control group showed the lowest expression level. Interestingly, cell stimulation in the PES ON group reduced HSP90 expression to 50% compared with the PES OFF group. Notably, this trend was consistent in both cell lines.

**Fig. 3.**
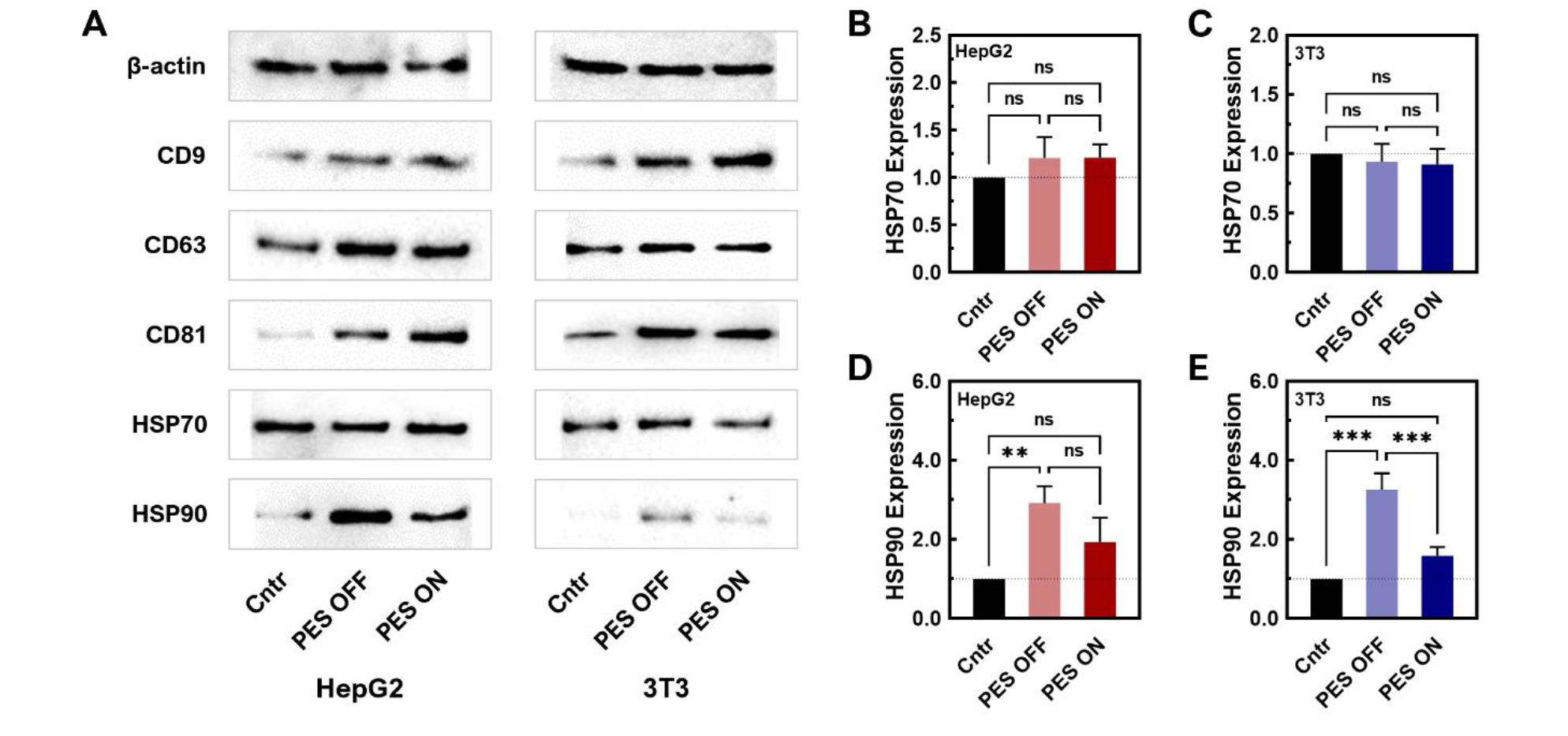
Characterization of sEVs in 2D culture control, PES OFF, and PES ON groups. **(A)** sEV marker characterization of CD9, CD63, and CD81, and HSP70 and HSP90 using western blot assays. **(B**–**C)** Quantitative measurement of HSP70 in **(B)** HepG2- and **(C)** 3T3-derived sEVs. Data shown as ± SEM of 3 replicate samples. **(D**–**E)** Quantitative measurement of HSP90 in **(D)** HepG2- and **(E)** 3T3-derived sEVs. Data shown as ± SEM of 3 replicate samples are representative of two independent experiments: \**P*<0.05, \*\**P*<0.01 \*\*\**P*<0.001, and \*\*\*\**P*<0.0001.

HSP70 and HSP90 have been reported as released from cells in EV involved in intercellular communication in cancer, immunity, and various pathological conditions.^59–61^ HSP70 is a molecular chaperone that helps in the proper folding of proteins. HSP70 in sEVs plays multifaceted roles in maintaining protein homeostasis, facilitating intercellular communication, modulating immune responses, and protecting cells from apoptosis. From our results, HSP70 levels did not change significantly, indicating the sEVs produced from 3D culture can maintain the protein cargo stability and homeostasis and thus fulfil its biological functions. HSP90 is an essential protein in protein folding, cancer progression and wound healing. It has been reported that HSP90 has been found to be a major cargo contained in sEVs. HSP90 in sEVs has several potential functions including selective client protein loading, stress response, and cancer progression and metastasis. However, the precise mechanisms and implications of HSP90’s role in sEVs are still being actively researched. In our study, we observed that HSP level increased significantly in PES OFF group while remained non-significant in PES ON group, compared to the control group. While HSP70 that stabilize protein did not change significantly and both non-cancer and cancer cells showed the similar trend, this observation indicated that cells in PES OFF group potentially produced more client protein loaded sEVs or displayed more stress response. Most importantly, PES ON group produced sEVs that contained similar level of HSP70 and HSP90 to control group, indicating minimum effect of acoustic stimulations on these two cargos.

In addition, we performed a cfDNA PCR analysis for genetic sEV cargo to understand the effect of the extracellular environment on the EV content. We tested two cargo sequences that code for key proteins in sEV biogenesis: NRAS, and P53 (*e*.*g*., Primers shown in **Table S1**). NRAS is a member of the ras-GTPase family that regulates proliferation and cell division by controlling the activation of the MAPK/ERK signaling pathway and PI3P pathways for cell growth, and cell survival.^62^ More importantly, NRAS is essential for sEV biogenesis, by mediating cargo selection and inducing sEV secretion.^63,64^ On the other hand, P53 is a transcription factor that functions as a tumor suppressing protein.^65^ In addition, previous reports have shown that sEV secretion can be promoted by the HSP-P53-TASP6 signalling pathway.^19,66^ We observed slight increases in genetic NRAS in HepG2 cell derived sEVs and no significant change in 3T3 derived sEVs (**Fig. S15A**). Meanwhile, upregulated TP53 were observed in genetic P53 content (**Fig. S15B**–**C**), suggesting that the 3D environment can affect key players in sEV biogenesis and promote pathways for induced sEV secretion. While this would support our findings of enhanced sEV production, further upregulation of cell proliferation pathways such as the MAPK/ERK signaling pathway and PI3P pathway can promote autophagy^67^, and tumorigenesis^68^, requiring future studies to understand the cell’s phenotypical characteristics under stimulated conditions. Most importantly, there is no significant difference between PES ON and PEF OFF samples, indicating that stimulated conditions did not induce unexpected changes in the sEV product.

### 3.3 Enhanced sEV production in stimulated PESs via increased cellular metabolic rate and intracellular calcium concentration

Understanding the factors that influence sEV biosynthesis and production in stimulative PESs is important for engineering suitable platforms for future applications. Previous studies have confirmed the impact of metabolic rate and increased metabolite levels on EV production, as several critical steps in sEV production rely on these metabolites for activation (**Fig. 4A**).^69^ Metabolites such as NADH and ATP play a pivotal role in promoting the ATP-dependent trafficking of multivesicular bodies (MVBs) for sEV secretion, thereby influencing sEV biogenesis and production. To investigate the underlaying mechanism of enhanced sEV production in the PES 3D culture platform, we conducted colorimetric assays measuring NADH hydrolysis activity (**Fig. 4B**) and ATP levels (**Fig. 4C**). Both NADH and ATP assays revealed increased activity in PES 3D culture systems with (PES ON) and without stimulation (PES OFF) compared with the traditional 2D culture (control), which is consistent with previous studies.^70,71^ Notably, we observed 24% increase in NADH hydrolysis activity and 16% increase in ATP levels in the PES OFF group compared with 2D culture control. Furthermore, we observed a 45% increase in NADH activity and ATP levels in the PES ON group compared with 2D culture control. This heightened metabolic activity in the stimulated 3D culture, for both representative non-cancer and cancer cells (*e*.*g*., 3T3 and HepG2), potentially stems from the increase in secondary messengers such as calcium ions, thus facilitating increased metabolism and sEV biogenesis and production.^72^

**Fig. 4.**
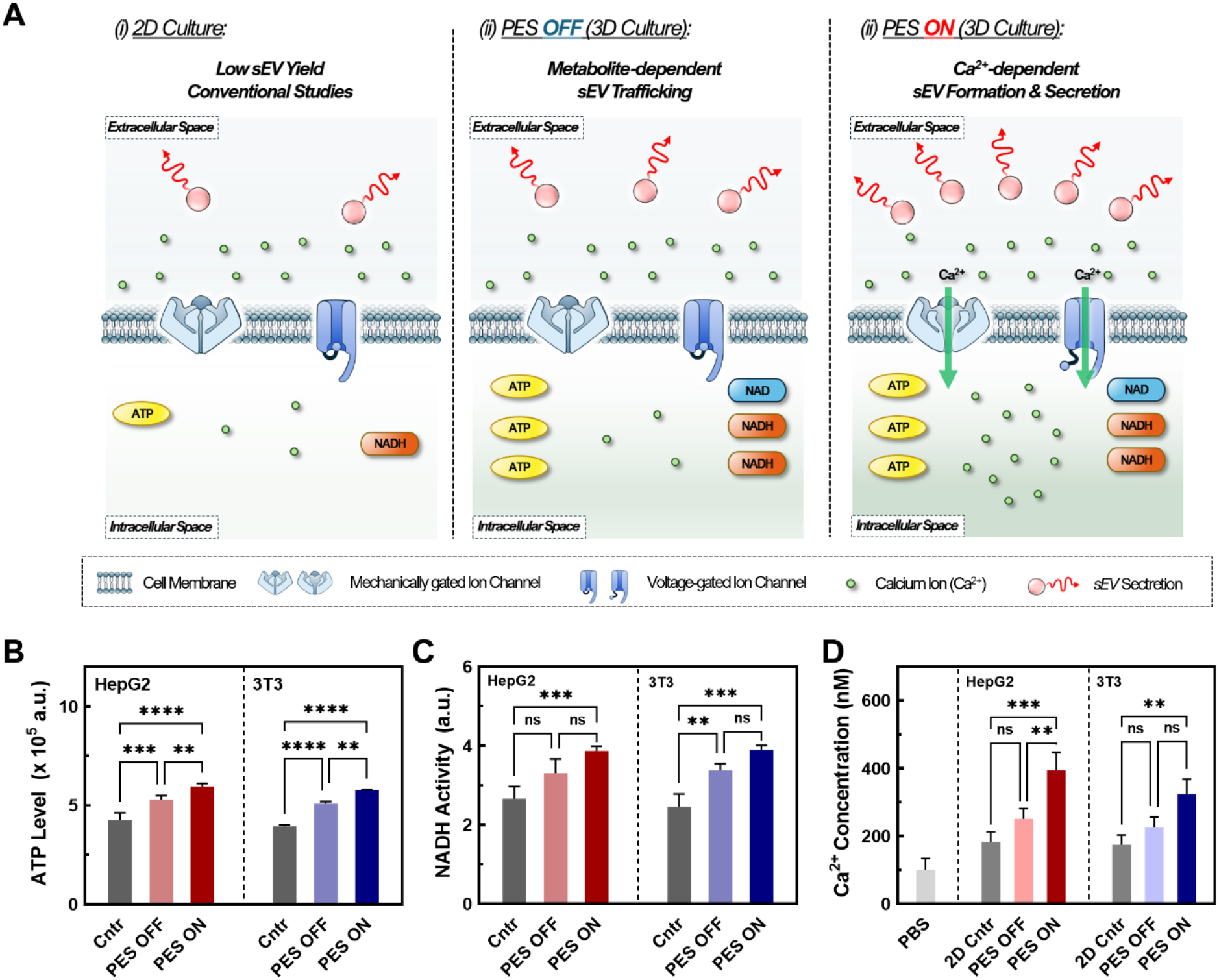
**(A)** Schematic of mechanisms for enhanced EVs through control, PES OFF, and PES ON groups. **(B)** ATP levels for all groups at various cell counts using Celltiter-glo. Data shown as ± SD of 3 replicate samples are representative of two independent experiments: \**P*<0.05, \*\**P*<0.01, \*\*\**P*<0.001, and \*\*\*\**P*<0.0001. **(C)** Comparison of NADH activity for all groups at various cell counts using CCK-8 assay. Data shown as ± SD of 3 replicate samples are representative of two independent experiments: One-way ANOVA analysis with \**P*<0.05, \*\**P*<0.01, and \*\*\**P*<0.001. **(D)** Relative Ca^2+^ concentrations of all groups measured by Fura-2AM. Data shown as ± SD of 3 replicate samples are representative of two independent experiments: \**P*<0.05, \*\**P*<0.01, and \*\*\**P*<0.001.

Previous studies have also highlighted the crucial role of elevated intracellular calcium in sEV production by triggering essential steps such as sEV formation, cargo recruitment, and sEV secretion.^13,19,25^ To investigate the potential role of calcium ions in our PES culture platform, we conducted intracellular calcium measurements using Fura-2AM (**Fig. 4D**). Across all groups, we noted a 1.5-fold increase in the intracellular calcium ion concentration in the PES ON group compared with the PES OFF group and control 2D culture, confirming its impact on sEV production. Various mechanisms account for this calcium influx. While electrical stimulation has been shown to promote cell membrane reorganization, leading to calcium influx,^18,19,25^ our system operates at 10–100-fold lower voltages to preserve cell viability and cell membrane integrity, suggesting the coexistence of alternative mechanisms. A possible pathway for calcium influx is the activation of voltage-gated calcium channels (VGCCs) such as L-, N-, and P-type VGCCs, which are opened through cell membrane depolarization requiring only 30 mV for activation.^52^ Low-frequency AC electric fields can also influence the cell membrane action potential and open VGCCs, allowing for an influx of Ca^2+^ ions into the cell.^73^ Alongside piezoelectric stimulation, audible acoustic stimulation can activate mechanosensing calcium channels such as Piezo-1 through mechanical force induction.^74,75^ Overall, we found that increased metabolite levels coupled with enhanced intracellular calcium levels potentially activated multiple stages of sEV production in both cell lines.

### 3.4 Impact of cell morphology in PES 3D culture platform on sEV production

While we observed a significant increase in sEV production yield and rate in the stimulated PES 3D culture platform, this synergistic effect varied across different cell lines. Therefore, we investigated other potential mechanisms that could have influenced sEV production in this platform. We first hypothesized that such a difference was attributable to the varied cell morphology in the 3D culture, with enhanced cell–matrix interactions, as well as in different cell types.^76,77^ This is based on recent studies showing that less well-developed actin cytoskeletons due to cell–matrix interactions, as reflected in cell morphology changes, promote intracellular MVB trafficking and resulting fusion with the plasma membrane.^78^ The inhibition of actin-related proteins restores MVB trafficking on the stiffer substrate, which leads to changes in EV production.^78^ In particular, when the cell morphology changes to a rounder shape, the cytoskeleton is redistributed to the plasma membrane, promoting MVB trafficking near the membrane for sEV secretion.^79^

To assess this hypothesis, we imaged both HepG2 and 3T3 cells in all groups using SEM (**Fig. 5A** and **Fig. S16**), and then analyzed cell morphology parameters including the roundness, cell axial ratio, and cell area (**Fig. 5B** and **Fig. S17**). With great biocompatibility (See **Fig. S6C**) and stable cell adhesiveness of our system (**Table S3**), morphological parameters were able to serve as indicators of membrane curvature and cell polarity.^78–80^ First, we compared these parameters between cells cultured in the 2D control and in 3D PES culture platform for each cell line. The results demonstrated that cell roundness in the PES OFF group is 1.3-fold higher (HepG2 cells) and 2.1-fold higher (3T3 cells) than that in the 2D culture. Interestingly, the cell roundness is highly correlated with the sEV production rate and the enhancement in production rate in PES (**Fig. 5C**), implying cell roundness under 3D culture as a key factor affected by cell type (See **Fig. 2D–E**). To quantify the correlation between sEV production and cell roundness, we fitted the data to the equation *Y = Y*_0_*e*^*kx*^, where *Y* is the sEV production rate or the enhancement in the production rate after stimulation, *Y*_0_ is an initial fitting parameter, *k* is the exponential fitting coefficient, and *x* is the cell roundness. Based on mathematical fitting, the cell roundness is exponentially correlated with the EV production rate with *k* = 3.33 ± 0.5 and is correlated with the enhancement in EV production in stimulated groups with *k* = 6.56 ± 0.8. These correlations indicate that cell roundness, influenced by the cell culture microenvironment in the 3D culture matrix, plays a key role in sEV production, and profoundly impacts the synergistic effect of EV production through both the 3D culture and cell stimulation. Additionally, by comparing the cell morphologies between cell lines, we further discovered that 3T3 cells consistently exhibited a 2.5-fold higher axial ratio than HepG2 cells across all groups (See **Fig. S17**). These results indicate that the more polarized cell morphology of 3T3 cells compared with HepG2 cells creates a more complicated sEV biogenesis pathway,^81–83^ potentially decreasing the sEV production rate in 3T3 cells compared with HepG2 cells.

**Fig. 5.**
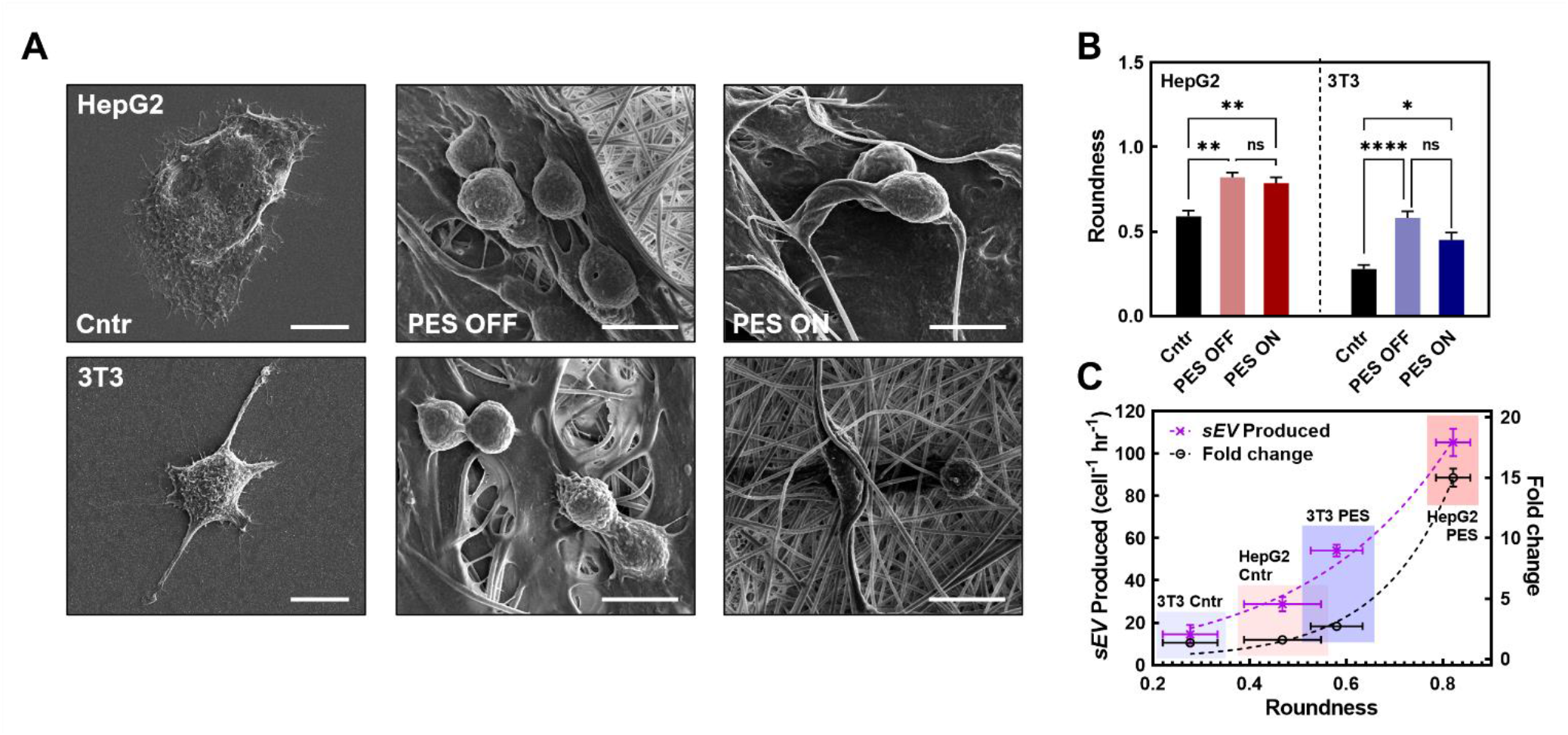
**(A)** SEM images of HepG2 and 3T3 fibroblast cells under three different cultures including 2D, PES OFF, and PES ON cultures. (Scale bar: 10 μm) **(B)** The cell measurement through roundness. **(C)** The correlation between the roundness and the sEV production rate and fold change in production rate post stimulation. All data shown as ± SD of 3 replicate samples is representative of two independent experiments: \**P*<0.05, \*\**P*<0.01 \*\*\**P*<0.001, and \*\*\*\**P*<0.0001.

In summary, we showcased the synergistic effect of employing electrical stimulation strategies for enhancing sEV production within a 3D culture model. This synergistic effect translated into 15.7-fold and 6.7-fold increases in sEV production rate compared with the traditional 2D culture for HepG2 and 3T3 cells, respectively. We postulated that this synergistic effect on sEV production was influenced by a 1.5-fold rise in intracellular calcium ions and a 40% increase in metabolite concentration. Furthermore, we identified the exponential correlation between cell roundness and the magnitude of sEV production enhancement post-stimulation, indicating that the cell morphology in the 3D culture platform based on PES plays a pivotal role in influencing sEV production efficiency.

Despite these findings, the specific interplaying mechanisms between factors such as Ca^2+^ remain unclear, requiring further investigation and understanding. Accordingly, optimization of scaffold parameters and stimulation conditions will be conducted for future applications to enable optimal sEV production. Furthermore, beyond the proof-of-concept studies, the use of human cell lines, including stem cells and white blood cells, will provide more clinically relevant therapeutic applications in future translational studies.

## 4. Conclusion

In this study, we systematically investigated the effect of 3D culture and cell stimulation on EV biomanufacturing. Our findings highlight the significance of this approach in the context of the scaled-up manufacture of EVs for clinical translation. Herein, it was demonstrated that the 3D stimulated culture produced EVs at a 15.7-fold increased rate per cell without any significant deviation in particle size or protein composition. Cells under the stimulated 3D culture also contained significantly higher metabolite and calcium ion concentrations, indicating that specific steps in sEV biogenesis are being activated and upregulated faster than in the standard 2D culture. These findings hold great promise for advancing EV therapeutics into clinical applications and overcoming the challenge of low EV production rates in standard methods. In addition, these findings in both non-cancer and cancer cells provide a promising 3D culture platform based on piezoelectric biomaterials for various applications, including investigation of the effect of bioelectricity on the metastatic behavior of cancer cells and the stimulated production of sEVs for advanced therapeutics in clinical settings.

## Author contributions

YW, JJ, and HC conceptualized the study. JJ and YW designed the experiments. JJ and YC synthesized, imaged, and characterized the PES under NM’s supervision. HJ imaged and characterized the cell culture in PES. JJ and CK performed cell-based experiments, and sEV isolation and concentration measurements. GK performed western blot experiments. TS performed qPCR analysis. YW, HJ, and JJ analyzed the data and wrote the original draft, and YW supervised all the work. All authors contributed to the article and approved the submitted version.

## Conflicts of interest

There are no conflicts to declare.

## Acknowledgements

This work was supported by the National Institutes of Health (NIH) under the Maximizing Investigators’ Research Award (MIRA) [R35GM150608], NIH National Cancer Institute [R21CA277663-01A1], and Berthiaume Institute for Precision Health (BIPH) under the Discovery Funding, and the University of Notre Dame under the Seed Transformative Interdisciplinary Research (STIR) Grant. All the cell-based SEM imaging, and TEM imaging were carried out in part in the Notre Dame Integrated Imaging Facility, University of Notre Dame, using the Magellan 400 field emission scanning electron microscope, and the Talos F200i field emission (scanning) transmission electron microscope. We thank Tatyana Orlova and Maksym Zhukovskyi for the knowledge and expertise as well as time towards this research. All NTA characterization was carried out in part in the Harper Cancer Research Institute Tissue Bank Facility using the Nanosight NS300.

## Notes

### Competing Interest Statement

The authors have declared no competing interest.

### Summary of Updates

we have undertaken extensive revisions throughout the manuscript, as follows: i) we conducted comprehensive characterization of the scaffolds, providing detailed description of their properties and functions; ii) we performed content analysis of the produced small extracellular vesicles (sEVs), and discussed their potential implications; iii) we investigated the underlying mechanisms responsible for the increased production of sEVs; iv) we thoroughly evaluated the performance of 3D cell cultures, focusing on critical parameters such as cell adhesion, penetration, and proliferation, allowing to present a more robust and detailed assessment of the 3D culture system efficacy; v) we reassessed the production rate of sEVs based on more accurate cell counts, ensuring the reliability and reproducibility of our results.

