## Supplemental information for "Stimulative piezoelectric nanofibrous scaffolds for enhanced small extracellular vesicle production in 3D cultures"

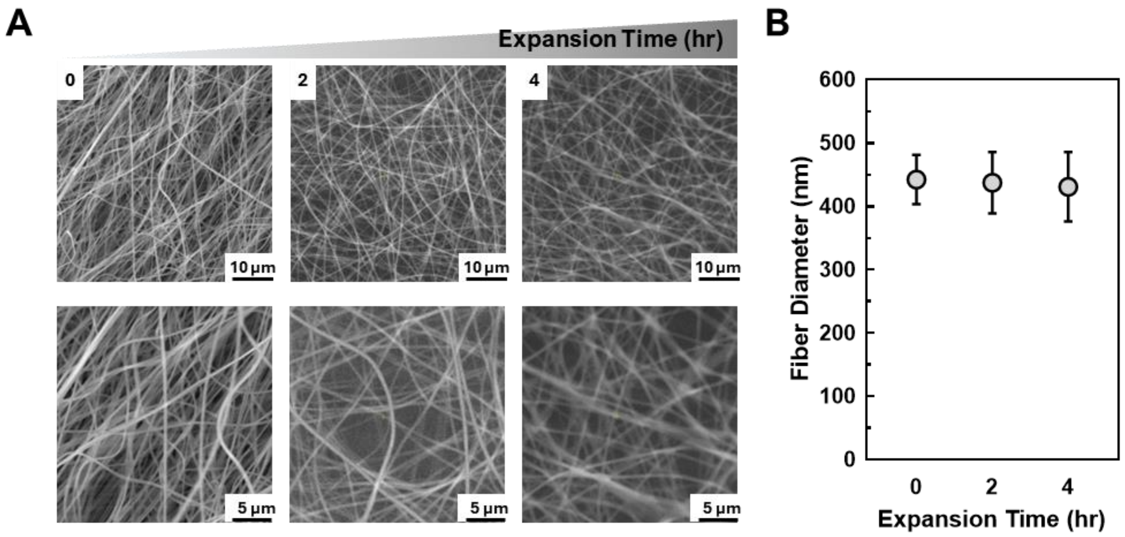


**Fig. S1. (A)** Scanning electron microscopy (SEM) images for the Piezoelectric scaffolds (PES) at 0, 2, and 4 hr of gas expansion time. **(B)** Measured fiber diameter inside PES along the gas expansion time.

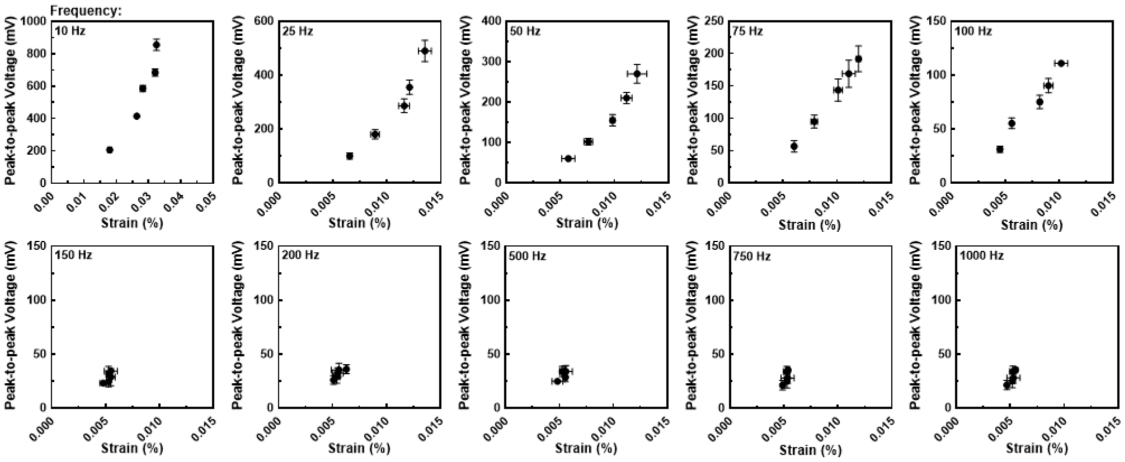


**Fig. S2.** The effect of acoustic amplitude on the piezoelectric properties of PES at frequencies of 10, 25, 50, 75, 100, 150, 200, 500, 750, and 1000 Hz. Scale bars are +/- 1 standard deviation (SD). (*N*=4).

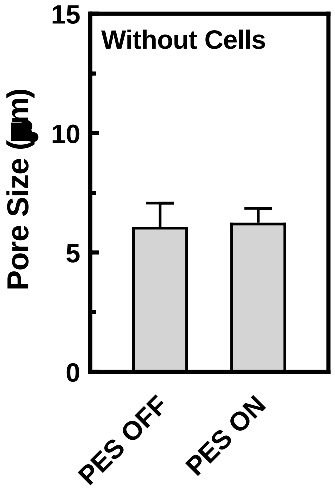


**Fig. S3.** The effect of acoustic stimulation on the pore size of PES. Scale bars are +/- 1 standard error of mean (SEM). ($N$=5).

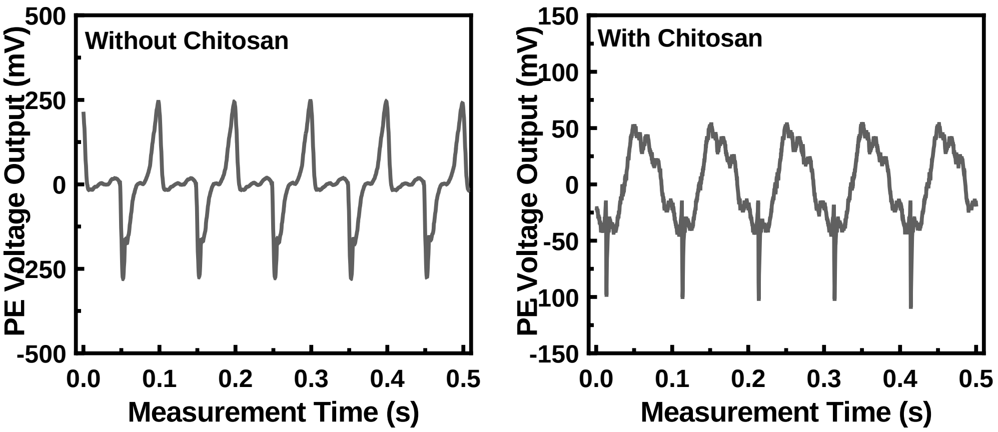


**Fig. S4.** PE output comparison between pristine scaffold (left) and chitosan-coated scaffold (right).

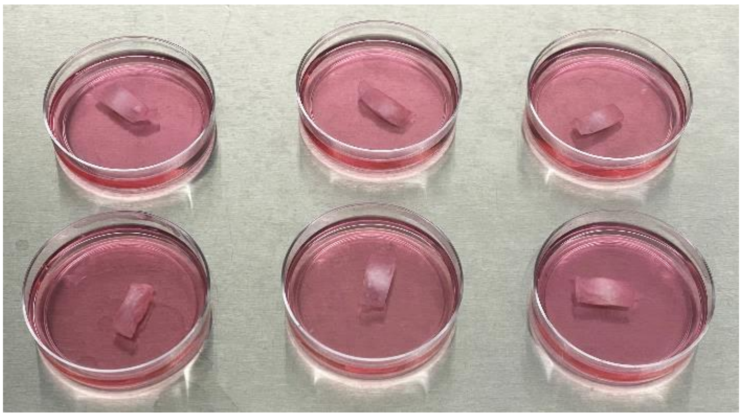

**Fig. S5**. Representative digital image of the scaffold inside the culture media.

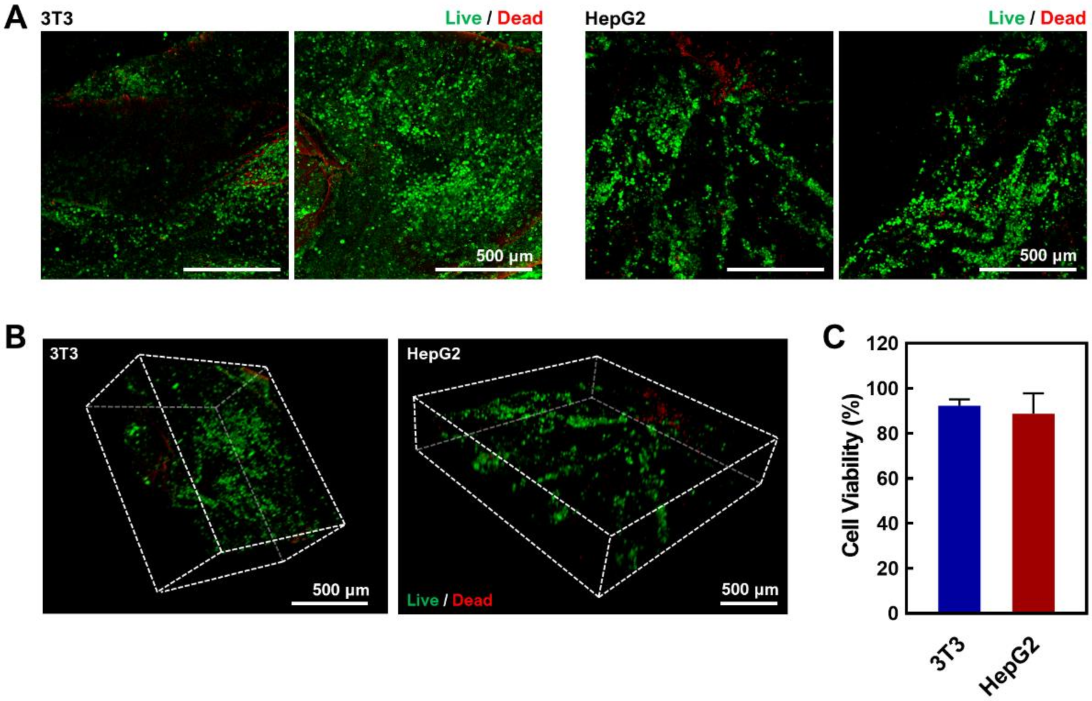


**Fig. S6**. Confocal light scanning microscopic images of 3T3 and HepG2 on chitosan-coated scaffolds stained with live/dead kit. **(A)** Maximum intensity projected images of both cell lines (Left: 3T3 and Right: HepG2). **(B)** 3D constructed confocal image of both cell lines (Left: 3T3 and Right: HepG2). **(C)** Cell viability comparison between 3T3 and HepG2. Red and green signal were collected from three confocal images for each cell lines (*N*=3).

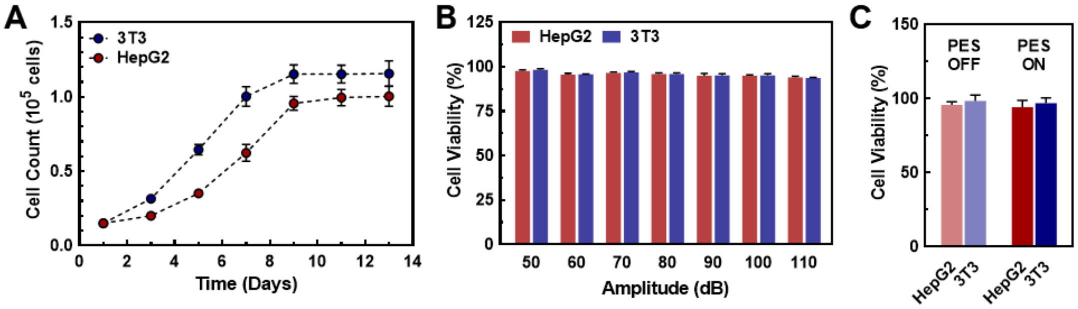


**Fig. S7**. The biocompatibility and cell proliferation on PES. **(A)** Cell proliferation after 14 days of cell culture on PES. **(B-C)** Cell viability after stimulation at amplitudes **(B)** between 50–110 dB and **(C)** 85 dB.

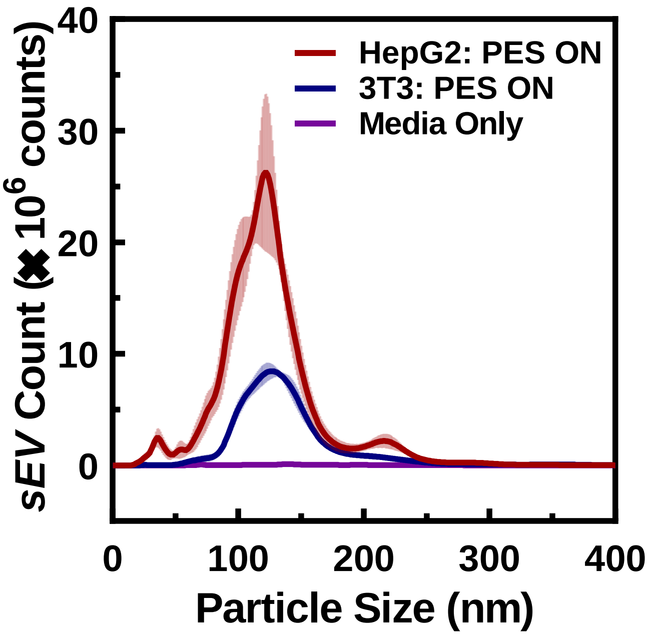


**Fig. S8**. Raw particle count from NTA analysis (*N*=5).

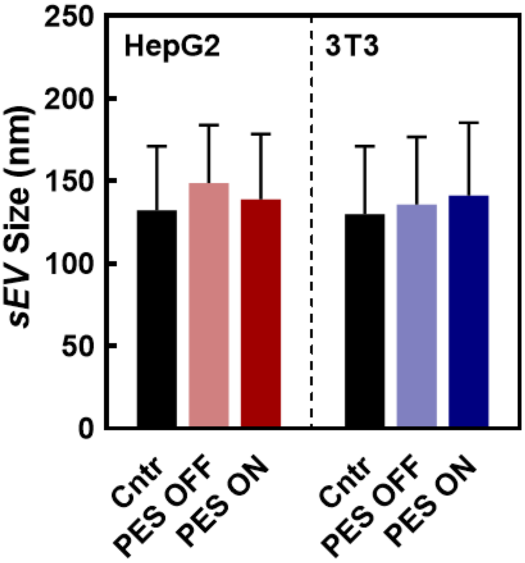


**Fig. S9**. sEV characterization using NTA. The mean particle size of the produced sEVs. Error bars are +/- 1 SD (*N*=5).

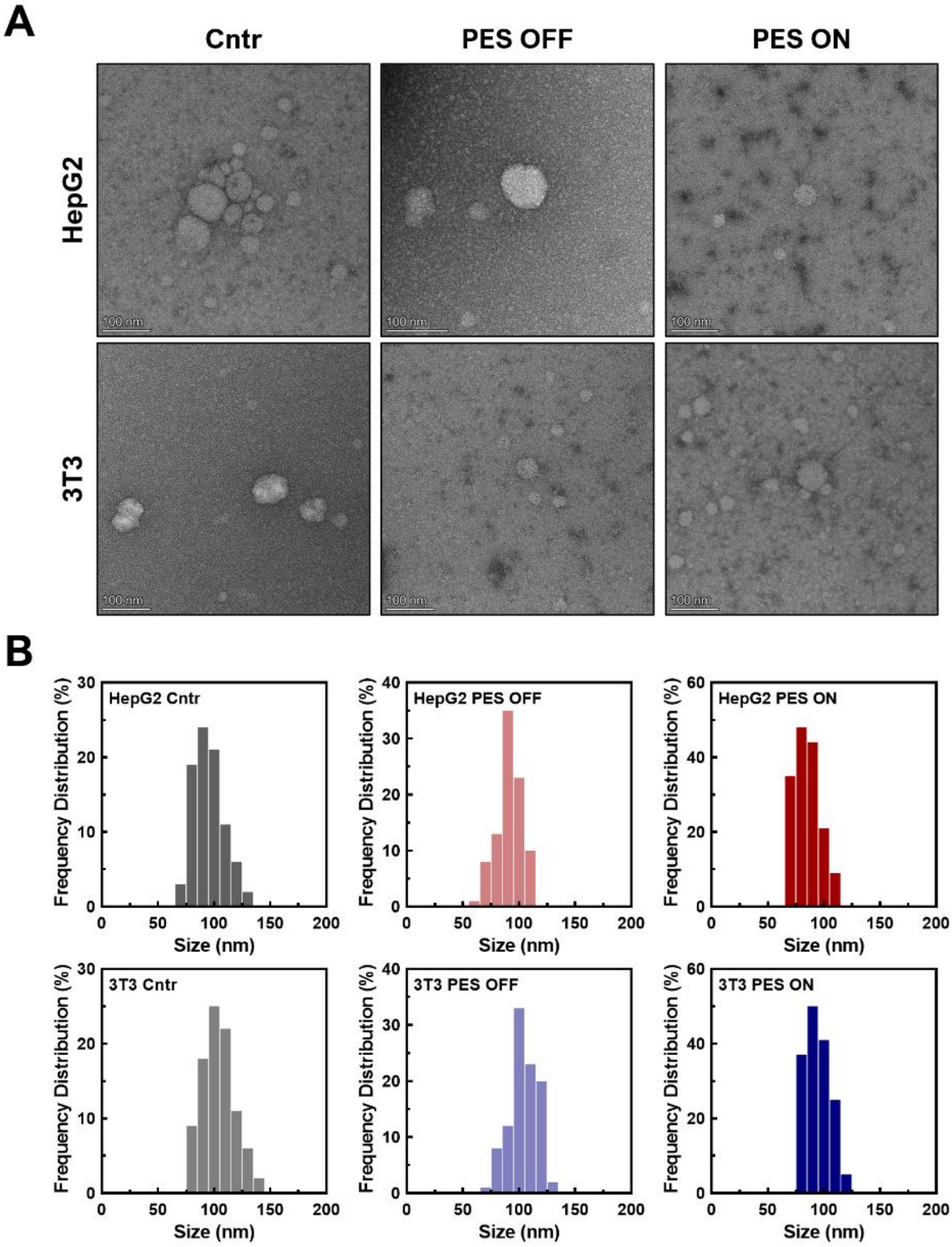


**Fig. S10.** Size distribution of sEVs through TEM. **(A)** TEM images of sEVs from control, and PES culture. **(B)** Size distributions histograms of sEVs based on TEM imaging.

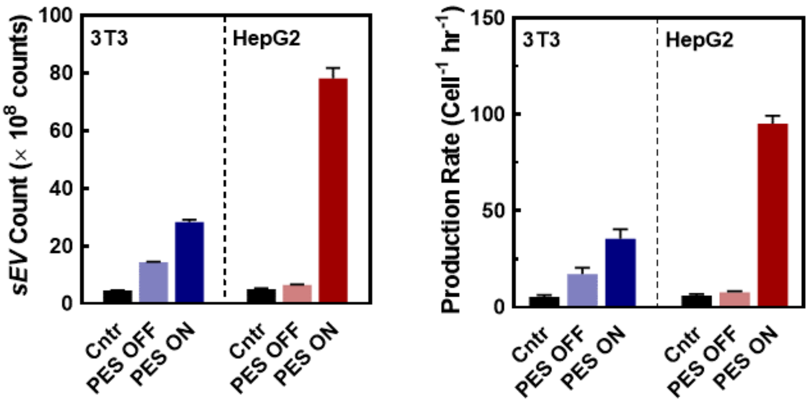


**Fig. S11.** The sEV Production measurements of PES with and without activation using NTA (SD; *N*=5).

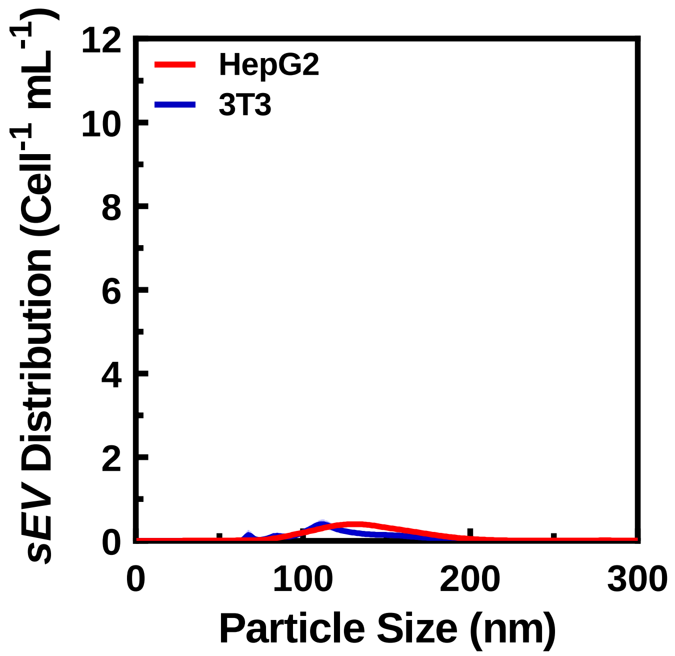


**Fig. S12**. The sEV production from control group with acoustic stimulation (Cntr ON; Red: HepG2, Blue: 3T3).

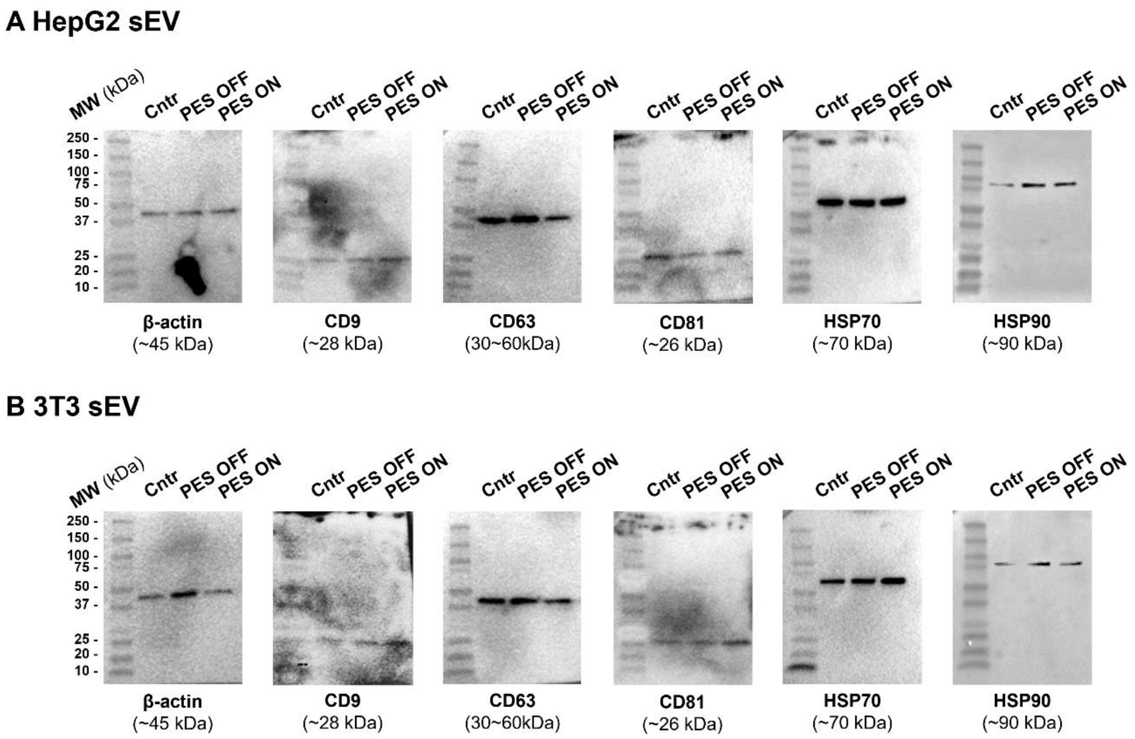


**Fig. S13**. Images of the complete western blot gels using the white light channel for **(A)** HepG2 derived sEVs and **(B)** 3T3 derived sEVs.

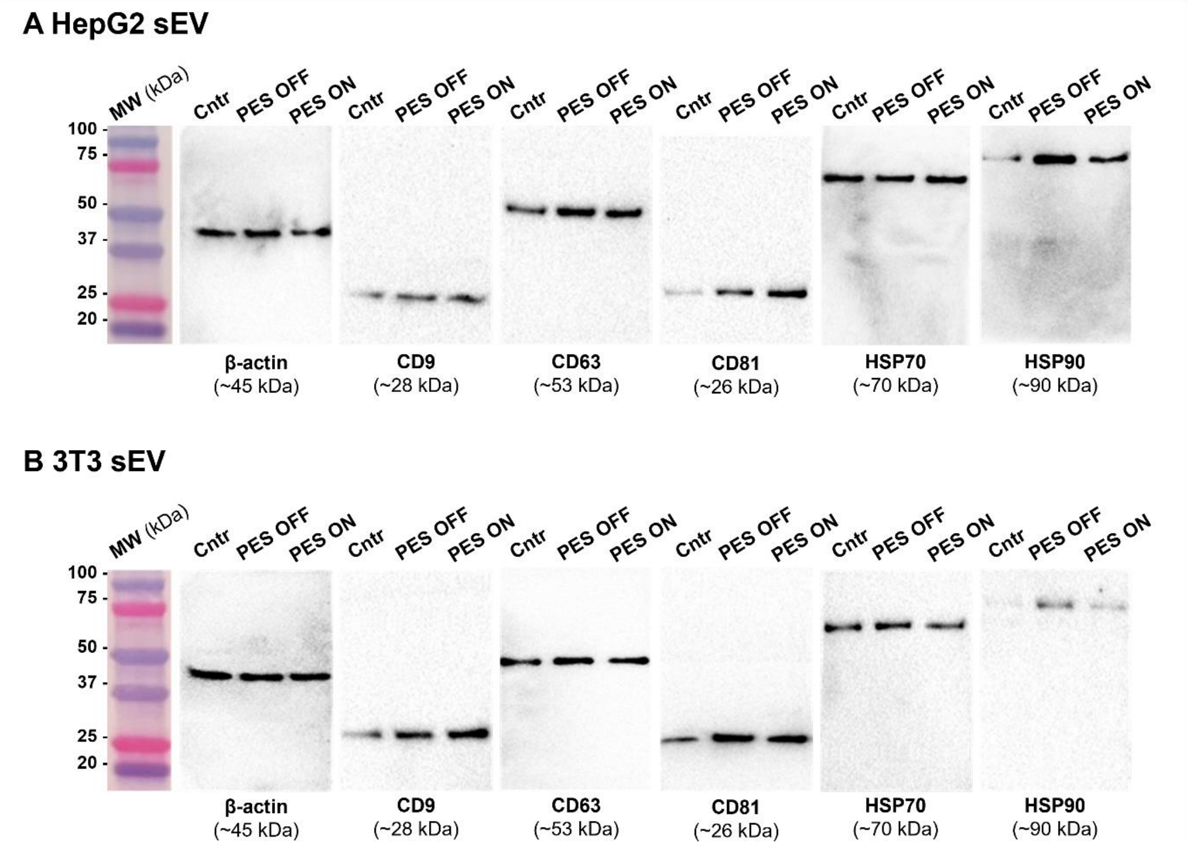


**Fig. S14**. Images of the western blot gels of **(A)** HepG2 derived sEVs and **(B)** 3T3 derived sEVs. The ladder was imaged using the color light channel, and the bands were imaged using the white light channel.

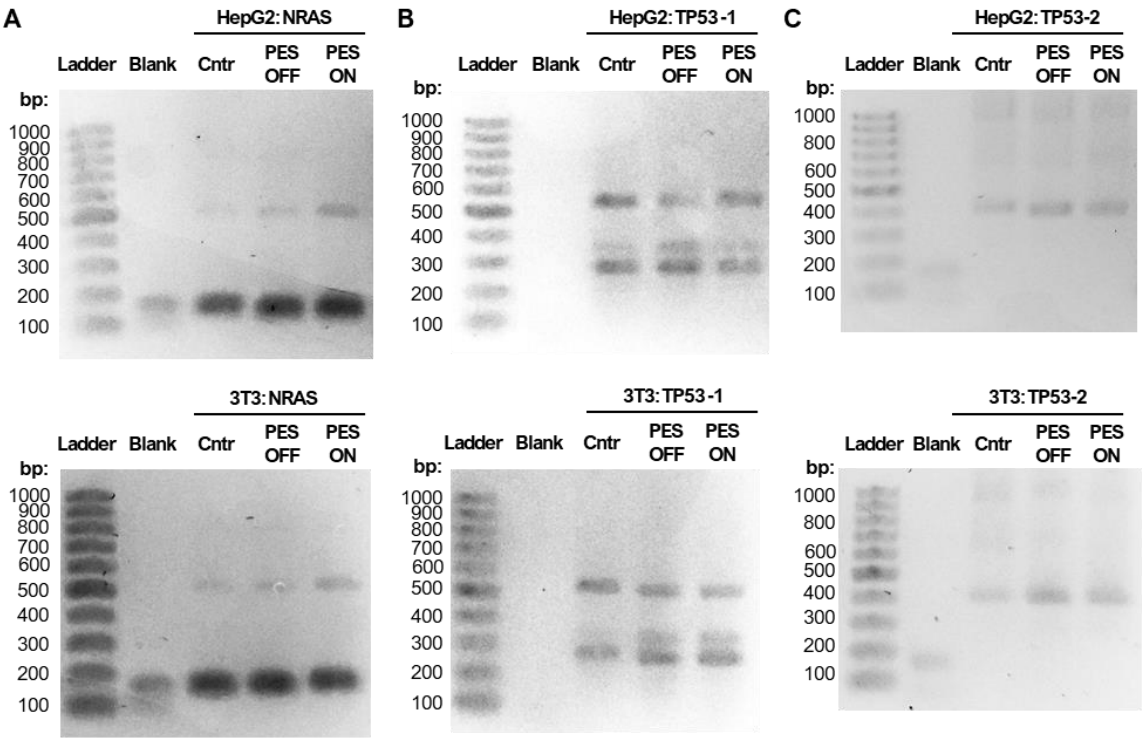


**Fig. S15.** Gel electrophoresis results from the cfDNA PCR of sEV content. **(A)** NRAS-targeting PCR result from HepG2 (Top) and 3T3 (Middle). Result from quantitative analysis for NRAS (Bottom). **(B)** TP53 Inner-targeting PCR result from HepG2 (Top) and 3T3 (Middle). Result from quantitative analysis for TP53-1 (Bottom). **(C)** TP53 Outer-targeting PCR result from HepG2 (Top) and 3T3 (Middle). Result from quantitative analysis for TP53-2 (Bottom).

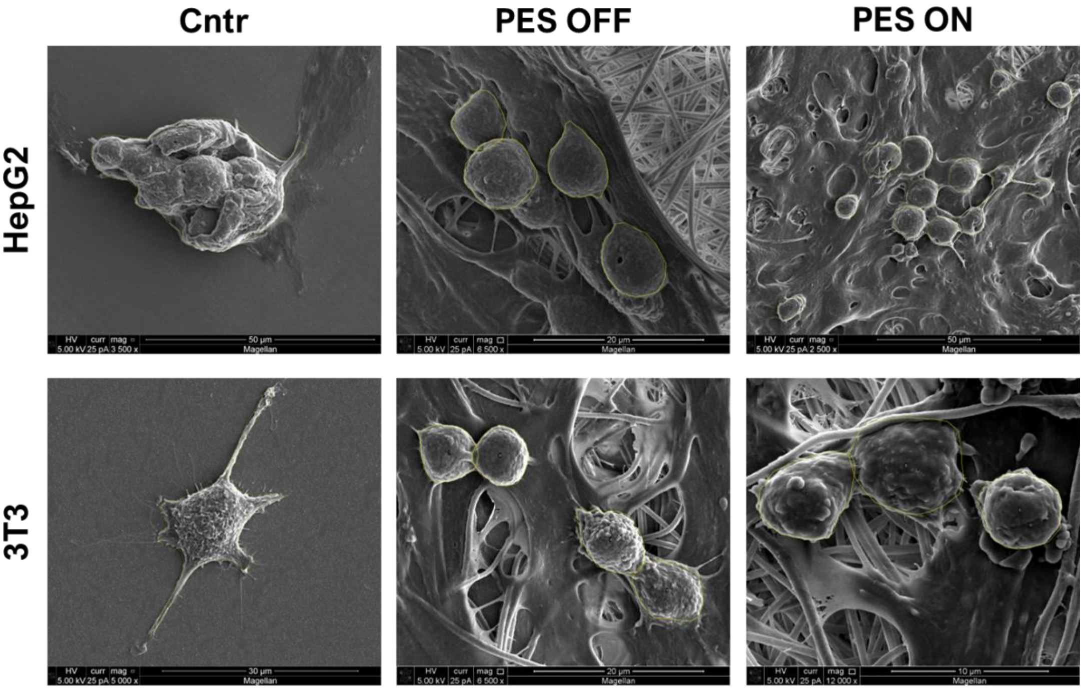


**Fig. S16.** Cell morphology analysis outlined SEM images.

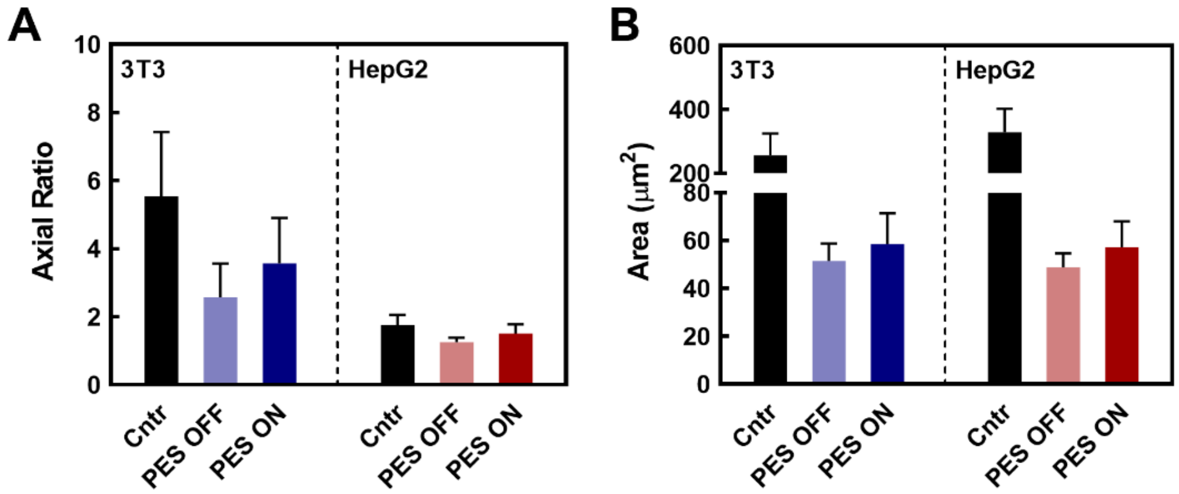


**Fig. S17.** Additional cell morphology measurement including **(A)** Axial ratio and **(B)** Cell area on 2D and PES culture platforms using SEM imaging. Scale bars are +/- 1 SD. (*N*=3).

**Table S1**. Primer set sequences for TP53 Nested PCR and NRAS PCR.

| PCR Reaction | Forward Primer (5’ → 3’) | Reverse Primer (5’ → 3’) |
|---|---|---|
| TP53 Outer (541 bp) | 5’-CTG AGT GAC AGA GCA AGA CCC TAT-3’ | 5’-AGT GTT TCT GTC ATC CAA ATA CTC C-3’ |
| TP53 Inner (397 bp) | 5’-GTT TCT TTG CTG CCG TCT TC-3’ | 5’-ACA CGC AAA TTT CCT TCC AC-3’ |
| NRAS (152 bp) | 5’-CAC AAA GAT CAT CCT TTC AGA GA-3’ | 5’-ACA AGA AGA GTA CAG TGC CA-3’ |

**Table S2**. Summary of characteristics and parameters of PAN solution.

| **Material / Solvent** | **Viscosity** (cP) | **Surface Tension** (mN/m) | **Electrical Conductivity** (µS/cm) | **Density** (g/mL) |
|---|---|---|---|---|
| 10%$_{wt}$ PAN in DMF | 411.8 | 27.5 | 38.5 | 1.194 |

**Table S3**. Result of cell adhesion-cell loss throughout cell seeding and stimuli processes.

| Cell Line / Platform | | Cell Loss (%) | | Cell Adhesion (%) |
|---|---|---|---|---|
| | | In Seeding | In Stimuli | |
| HepG2 | **2D Non-adhesive** | CCK: **90.9 ± 9.21**<br>PB: **91.0 ± 9.65** | CCK: 3.95 ± 1.78<br>PB: 3.80 ± 2.17 | CCK: 5.13 ± 1.26<br>PB: 5.24 ± 1.15 |
| | **2D Adhesive** | CCK: 1.78 ± 0.45<br>PB: 1.80 ± 0.38 | CCK: 0.67 ± 0.31<br>PB: 0.64 ± 0.37 | CCK: **97.6 ± 0.85**<br>PB: **97.6 ± 0.86** |
| | **3D PES OFF** | CCK: 1.88 ± 0.15<br>PB: 1.87 ± 0.35 | CCK: 1.35 ± 0.38<br>PB: 1.31 ± 0.25 | CCK: **96.8 ± 1.08**<br>PB: **96.8 ± 1.10** |
| | **3D PES ON** | CCK: 1.73 ± 0.16<br>PB: 1.67 ± 0.08 | CCK: 1.00 ± 0.23<br>PB: 1.04 ± 0.25 | CCK: **97.3 ± 2.44**<br>PB: **97.3 ± 2.44** |
| 3T3 | **2D Non-adhesive** | CCK: **76.7 ± 6.30**<br>PB: **91.0 ± 9.65** | CCK: 8.27 ± 0.37<br>PB: 3.80 ± 2.17 | CCK: 15.0 ± 2.24<br>PB: 5.24 ± 1.15 |
| | **2D Adhesive** | CCK: 1.43 ± 0.16<br>PB: 1.8 ± 0.38 | CCK: 0.72 ± 0.04<br>PB: 0.64 ± 0.37 | CCK: **97.9 ± 0.81**<br>PB: **97.6 ± 0.86** |
| | **3D PES OFF** | CCK: 1.34 ± 0.14<br>PB: 1.87 ± 0.35 | CCK: 0.58 ± 0.07<br>PB: 1.31 ± 0.25 | CCK: **98.1 ± 0.99**<br>PB: **96.8 ± 1.10** |
| | **3D PES ON** | CCK: 1.34 ± 0.18<br>PB: 1.67 ± 0.08 | CCK: 0.75 ± 0.18<br>PB: 1.04 ± 0.25 | CCK: **97.9 ± 1.20**<br>PB: **97.3 ± 2.44** |
